# Semaphorin3F Drives Dendritic Spine Pruning through Rho-GTPase Signaling

**DOI:** 10.1101/2021.03.05.433425

**Authors:** Bryce W. Duncan, Vishwa Mohan, Sarah D. Wade, Young Truong, Alexander Kampov-Polevoi, Brenda R. Temple, Patricia F. Maness

## Abstract

Dendritic spines of cortical pyramidal neurons are initially overproduced then remodeled substantially in the adolescent brain to achieve appropriate excitatory balance in mature circuits. Here we investigated the molecular mechanism of developmental spine pruning by Semaphorin 3F (Sema3F) and its holoreceptor complex, which consists of immunoglobulin-class adhesion molecule NrCAM, Neuropilin-2 (Npn2), and PlexinA3 (PlexA3) signaling subunits. Structure-function studies of the NrCAM-Npn2 interface showed that NrCAM stabilizes binding between Npn2 and PlexA3 necessary for Sema3F-induced spine pruning. Using a mouse neuronal culture system, we identified a dual signaling pathway for Sema3F-induced pruning, which involves activation of Tiam1-Rac1-PAK1-3 -LIMK1/2-Cofilin1 and RhoA-ROCK1/2-Myosin II in dendritic spines. Inhibitors of actin remodeling impaired spine collapse in the cortical neurons. Elucidation of these pathways expands our understanding of critical events that sculpt neuronal networks and may provide insight into how interruptions to these pathways could lead to spine dysgenesis in diseases such as autism, bipolar disorder, and schizophrenia.

## INTRODUCTION

During critical periods of cortical development there is an initial overproduction of dendritic spines and excitatory synapses on pyramidal neurons followed by activity-dependent elimination during the juvenile to adult transition [1]. Whereas much is known about activity-dependent spine elimination in the adult brain [2], much less is understood about developmental spine pruning. Identification of postnatal mechanisms for regulating spine and synapse density is important for defining normal circuit refinement, as well as providing mechanistic insight into how spine density is altered in neurodevelopmental disorders such as autism, schizophrenia, and bipolar disorder [3].

L1 family cell adhesion molecules (L1, NrCAM, Close Homology of L1 (CHL1), Neurofascin) play diverse and essential roles in the developing brain by promoting axon growth, cell adhesion, and migration [4]. L1-CAMs also mediate repellent axon guidance in response to class 3 Semaphorins [5–7], which are secreted ligands that stimulate growth cone collapse and axon repulsion [8]. Recently it was found that Sema3F and Sema3B induce retraction and loss of dendritic spines in postnatally developing cortical pyramidal neurons [9–11]. These ligands display striking selectivity in pruning different populations of spines even on the same dendrite. This selective spine pruning is achieved through a combinatorial mechanism in which Sema3 holoreceptor complexes comprising L1-CAMs, Neuropilins (Npn1/2) and PlexinA subunits (PlexA1-4) associate in different combinations to transduce intracellular signaling leading to spine collapse. Specifically, NrCAM and CHL1 function as obligate subunits of the Sema3F and Sema3B receptor complexes, respectively. Both L1CAMs constitutively bind Npn2, which can recruit different PlexA subunits [10,12,11]. The NrCAM, Npn2, PlexA3 complex mediates Sema3F-induced spine pruning [10], whereas the CHL1, Npn2, PlexA4 complex mediates Sema3B-induced spine pruning [12]. Single null mutant mice lacking NrCAM, CHL1, Npn2, PlexA3, or Sema3F display elevated spine and synapse density only on apical dendrites, where Npn2 is preferentially localized [9]. Selective spine pruning on apical dendrites is of interest with regard to cortical network development, as apical dendrites differ from basal dendrites in having more robust intra-cortical and thalamo-cortical inputs, and distinct temporal plasticity rules [13, 14]. Sema3B [12] and Sema3F [15] are secreted in an activity-dependent manner, raising the possibility that limited diffusion of Sema3 ligands may prune inactive neighbors during refinement of cortical circuits.

The specific role played by L1-CAMs within Sema3 receptor complexes and downstream Sema3 signaling mechanisms that bring about selective spine pruning are not well defined. To address these issues, we focused on the Sema3F signaling complex comprising NrCAM, Npn2, and PlexA3 (Fig. 1 A). The NrCAM extracellular region has 6 immunoglobulin (Ig) domains and 5 fibronectin III repeats, while the Npn2 ectodomain consists of 2 CUB domains (a1, a2), 2 coagulation factor V/VII domains (b1,b2) and a meprin-A5-mu domain (MAM). NrCAM binds Npn2 at an interface between the NrCAM Ig1 and Npn2 a1 domains in the absence of PlexA3 or Sema3F [10, 12]. Neuropilins bind PlexinAs with modest affinity, which is increased upon binding of dimerized Sema3 ligands [16]. Binding of Sema3 dimers to holoreceptor complexes stimulates the intrinsic Rap1-GTPase activating protein (GAP) activity of PlexA subunits through receptor dimerization and conformational change [17]. NrCAM induces clustering of Npn2 and PlexA3 in the neuronal membrane, stimulating PlexA3 Rap-GAP activity, which in turn downregulates Rap1-GTPase and inactivates β1 integrins [12]. Loss of integrin-dependent adhesion to the extracellular matrix may be an early step in spine pruning, since activated β1 integrins promote spine stabilization [18]. However, the downstream signaling events that cause spines to retract have not been elucidated.

**Fig. 1.**
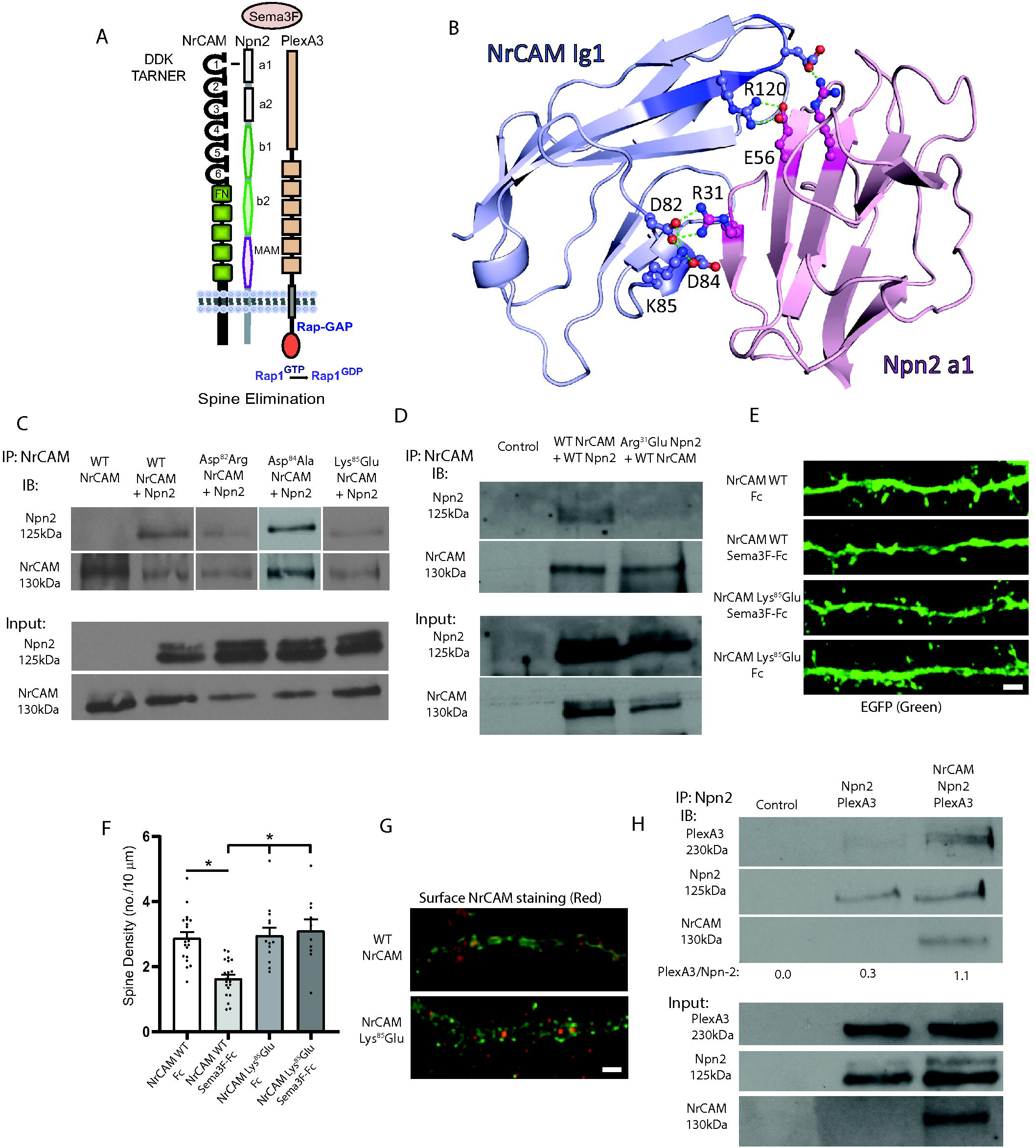
Structural Interactions of NrCAM in the Sema3F Holoreceptor Complex. A. Domain structure of the Sema3F holoreceptor complex comprising NrCAM (Ig1-6 and fibronectin III repeats), Npn2 (a1, a2, b1, b2 and MAM) domains, and PlexA3 with intrinsic Rap-GAP activity. B. Structural model for NrCAM Ig1 (blue) binding to Npn2 a1 (pink) Critical residues in the domain are rendered in ball-and-stick. Residue R^120^ in NrCAM TAR^120^NER motif is shown interacting with Npn2 E56. Residues D82, D84, and K85 in NrCAM Ig1 represent the “DDK” network, which is predicted to interact with R31 in the Npn2 a1 domain. The residue numbering represents the mouse sequences. C. Co-immunoprecipitation of WT NrCAM and NrCAM mutants Asp^82^Arg, Asp^84^Ala, and Lys^85^Glu with WT Npn2 from transfected HEK293T cells (equal amounts of extract protein), shown by immunoprecipitation (IP) of NrCAM and immunoblotting for Npn2 (IB). Blots were reprobed for NrCAM (lower panels). Inputs show equivalent levels of Npn2 and NrCAM in cell extracts. The blots shown are representative of 3 experiments. D. Co-immunoprecipitation of WT NrCAM with either WT Npn2 or Npn2 mutant Arg^31^Glu from transfected HEK293T cells (equal amounts extract protein). Blots were reprobed for NrCAM (lower panels). Equivalent levels of NrCAM and Npn2 were detected in input blots. The blots shown are representative of 3 experiments. E. Representative images showing NrCAM null neurons transfected with NrCAM Lys^85^Glu mutant or WT NrCAM plasmids in pCAGG-IRES-EGFP, treated with Fc or Sema3F-Fc for 30 min, immunostained for EGFP, and apical dendrites imaged confocally. Scale bar = 2µm. F. Quantification of mean spine density ± SEM on apical dendrites (n > 500 spines from ≥ 11 images per condition). Each point represents the mean spine density per image. 2-way ANOVA with Tukey post-hoc pairwise comparisons showed statistical significance (Control Fc vs. Sema3F-Fc, p = 0.021; Control Sema3F-Fc vs. K91E Fc, p = 0.008; Control Sema3F-Fc vs. K91E Sema3F-Fc, p = 0.004). G. NrCAM immunofluorescence staining (red) of unpermeabilized cortical neurons in cultures from NrCAM null mice transfected with WT NrCAM or NrCAM Lys^85^Glu plasmids. Plasmids included EGFP reporter to allow visualization of spines. Scale bar = 2µm. H. HEK293T cells were transfected with Npn2 and PlexA3 with or without NrCAM and cell lysates were assayed using co-immunoprecipitation (IP) of Npn2. Immunoblotting (IB) showed greater levels of PlexA3 in Npn2 IPs in the presence of WT NrCAM. Input blots show equivalent protein levels. The blots and quantification shown are representative of 3 experiments.

To understand the role of NrCAM within the Sema3F holoreceptor and to identify the critical signal transduction pathways leading to spine elimination, we carried out a structure-function study to identify key determinants in the NrCAM-Npn2 interface, then probed for downstream signaling effectors in a cortical neuron culture system. Here we identify a network of charged residues at the NrCAM-Npn2 interface that mediates association of NrCAM and Npn2, enhances Npn2-PlexA3 binding, and promotes Sema3F-induced spine pruning. We further show that Sema3F triggers a dual signaling cascade in cortical neurons regulated by small GTPases Rac1 (Rac1-PAK1-3 -LIMK1/2-Cofilin1) and RhoA (RhoA-ROCK1/2-Myosin II), resulting in rapid loss of dendritic spines achieved through actin cytoskeletal remodeling.

## MATERIALS AND METHODS

### NrCAM and Npn2 Structural Modeling, Mutagenesis, and Co-Immunoprecipitation

Structural modeling of the interaction between NrCAM Ig1 and Npn2 a1 domains was previously described [12]. Briefly, the extracellular NrCAM Ig1-Ig4 domains were subjected to homology modeling with MODELLER [19], based the crystal structure of human Neurofascin (PDB ID 3P3Y) [20]. Human Neurofascin and NrCAM are closely related homologs that share 49% overall sequence identity and 63% identity within the Ig1 domain. Npn2 a1-b2 domains were modeled based on the structure of Npn1 a1-b2 (PDB 4GZ9) [16] also using MODELLER. Mouse Npn2 and Npn1 share 63% amino acid homology [21]. The ClusPro protein-protein docking server was used for protein-protein interaction prediction between the NrCAM Ig1 and Npn2 a1 domains [22, 23]. The modeling was based on human sequences, while mutagenesis probed candidate interfaces in the mouse proteins. For NrCAM Ig1 the human and mouse sequences differ at only 2 out 101 residues, while for Npn2 a1 they differ at 2 out of 116 residues with one conservative substitution (Arg for Lys) near the predicted interface.

NrCAM Ig1 and Npn2 a1 mutations at the predicted NrCAM-Npn2 binding interface were generated using Q5 Site-Directed Mutagenesis (New England Biolabs, Ipswich, MA). Mouse NrCAM splice variant 1 (GenBank accession number AJ543321) in pCMV6 and mouse Npn2 (accession number NM_001077404) with an N-terminal FLAG tag in pcDNA3.1 were targeted for mutagenesis. Protein-protein interactions were assessed by co-immunoprecipitation from transfected HEK293T cells. HEK293T cells (from Bryan Roth, UNC) were grown in Dulbecco’s Modified Eagle’s medium-H supplemented with gentamicin/kanamycin and 10% fetal bovine serum (Corning #35-015-CV). Cells were seeded at 2 x 10^6^ cells/ 100mm dish the day before transfection. Plasmids were transfected with Lipofectamine 2000 (Invitrogen #11668030) in Opti-MEM. Media was changed to complete DMEM after 18 hours, and cells were lysed and collected 48 hours post-transfection. Cell lysates were prepared in Brij98 lysis buffer (1% Brij98, 10 mM Tris-Cl pH 7.0, 150 mM NaCl, 1mM EDTA, 1mM EGTA, 200 µM Na3VO4, 10 mM NaF, protease inhibitors (SigmaAldrich #P8340). Lysates (0.5–1 mg) were pre-cleared for 30 min at 4°C using Protein A/G Sepharose. Pre-cleared lysates (equal amounts of protein) were incubated with rabbit polyclonal antibody to NrCAM (Abcam #24344, RRID:AB_448024) or nonimmune IgG (nIg) for 2 h at 4°C. Protein A/G Sepharose beads were added for 30 min at 4°C prior to washing, and bound proteins were eluted by boiling in SDS-PAGE sample buffer. Western blotting was carried out using mouse monoclonal antibody clone M2 directed against the FLAG-tag on Npn2 (SigmaAldrich #F3165) or rabbit anti-NrCAM antibody (R&D Systems AF8538) and Avansta enhanced chemiluminescence. Protein bands were exposed within the linear response range on Xray films and quantified by densitometry to yield pixel densities. The ratio of mutant Npn2/NrCAM in the immunoprecipitate was normalized to the ratio of WT Npn2/WT NrCAM immunoprecipitated. Normalized ratios from 3 replicate experiments for each mutant were averaged and reported ± SEM.

To evaluate the role of NrCAM in stabilizing the association between Npn2 and PlexA3, HEK293T cells were transfected with pcDNA3.1-Npn2-FLAG and pcDNA3-mouse PlexA3, together with pCMV6-NrCAM or empty pCMV6 vector. After 48 hr Sema3F-Fc (5 nM) was added to the cultures for 20 min at 37°C. Cells were transferred to 4°C, lysed in Brij-98 lysis buffer, and clarified by centrifugation. Equal amounts of protein lysates were subjected to immunoprecipitation of Npn2 complexes using anti-Npn2 antibodies (R&D # AF567) and Protein A/G-Sepharose. Western blotting was performed using antibodies to PlexA3 (R&D #AF4075), Npn2 (FLAG tag) (SigmaAldrich,clone M2, #F3165), and NrCAM (R&D #8538). Protein bands were exposed within the linear response range on Xray films and quantifed by densitometry. In all studies protein concentration was determined by the bicinchoninic acid assay.

### Sema3F-induced Spine Retraction Assay in Mouse Cortical Neuron Cultures

Wild type (WT) C57Bl/6J mice (RRID:IMSR_JAX:000664) mice were maintained according to the University of North Carolina Institutional Animal Care and Use Committee (IACUC) policies (AAALAC Institutional Number: #329) and in accordance with NIH guidelines. All animal procedures were approved by the UNC IACUC (IACUC ID# 18-073). Dissociated cortical neurons from WT E15.5 mouse embryos (C57Bl/6J) were plated onto poly-D-lysine and laminin-coated Lab-Tek II chamber slides (1.5 x 10^5^ cells/well) or T25 cell culture flasks (1.5 x 10^6^ cells/flask) as described [12]. At 11 days *in vitro* (DIV11), cells were transfected with pCAGG-IRES-mEGFP and/or other plasmids using Lipofectamine 2000. On DIV14 cells were treated with human Fc (Abcam #90285) or recombinant mouse Sema3F-Fc fusion protein (R&D Systems #3237-S3) (5 nM) for 2-30 min to induce spine retraction. Cultures were fixed and stained using 4% paraformaldehyde (PFA), quenched with 0.1M glycine, permeabilized with 0.1% Triton X-100, and blocked with 10% goat serum (Sigma #G6767). To enhance the EGFP signal immunofluorescence staining was performed using chicken anti-GFP (Abcam #13970, RRID:AB_300798) and AlexaFluor AF488-conjugated goat anti-chicken secondary antibodies (Thermo Fisher # A-11039, RRID:AB_2534096). For analysis of spine density, confocal imaging was performed on a Zeiss LSM 700 microscope in the UNC Microscopy Services Laboratory. As described in [12] approximately 10 images of apical dendrites of labeled neurons with pyramidal morphology were obtained for each condition. Z-stacks were obtained using 0.2 µm optical sections of field size 67 x 67 µm, captured using a 40 X oil objective with 2.4 x digital zoom. The same laser power and gain settings were used for all images to ensure consistent quantification. 3D deconvolution was carried out using AutoQuant version 3 software (Media Cybernetics) with default blind settings and Imaris (Bitplane) software. Spines were traced and scored blind to observer using Neurolucida software from confocal maximum intensity z-stacks on 30 µm segments of the first branch of apical dendrites. Mean spine densities (no./10 μM ± SEM) were calculated and compared by two-way ANOVA with Tukey posthoc testing for multiple comparisons with significance set at p < 0.05.

The effect of Rac1 and RhoA inhibition on spine density was assessed by pretreatment of WT neuronal cultures on DIV14 with Rac1 inhibitor NSC23766 (50 µM, 1 hr) or transfection with dominant negative (DN) mutants Rac1-17N or RhoA-G17A in pcDNA (provided by C. Der, UNC) together with pCAGG-IRES-mEGFP. For Tiam1/2-GEF inhibition GEF-dead Tiam1 mutant (Q^1191^K^1195->^AA) in pFlag-CMV2 or Tiam1 shRNA in pSuper (provided by K.Tolias, Baylor College of Medicine) [24, 25]. To inhibit PAK1-3 cells were preincubated with the small molecule pyridopyrimidinone inhibitor FRAX 486 (Tocris, Inc.) [26] at 500 nM in 0.01% DMSO for 30 min. FRAX 486 is selective for group I PAKs 1-3. In other studies neurons were preincubated with an inhibitor of Rho-associated, coiled-coil containing protein kinase (ROCK1/2) Y27632 (Tocris, Inc.) at 10 µM in PBS for 1-2 hr, or the LIMK1/2 inhibitor SR-7826 (Tocris, Inc.) at 1 µM in 0.01% DMSO for 30 min. For inhibition of actin remodeling, cells were preincubated with Jasplakinolide or Latrunculin A (ThermoFisher, Inc.) at 1 or 2 µM in 0.01% DMSO for 20 min.

### RhoA, Rac1, Cdc-42 Pulldown Assays

To assess the activation of small GTPases, neuronal cultures (DIV14) were treated with Fc or Sema3F-Fc (5 nM) for 2-30 min. GTP-bound Rac1, RhoA, or Cdc42 were assayed as described [27] with modifications. Cells were lysed in 50 mM Tris pH 7.6, 150 mM NaCl, 1% Triton X-100, 0.5 mM MgCl2, 200 µM Na3VO4, and protease inhibitors. GTP-bound, activated Rac1 or Cdc42 were pulled down from cell lysates using an immobilized GST fusion protein of the PAK binding domain (PDB) of murine p65 PAK, which binds only the GTP-loaded molecules. To assay RhoA-GTP, a GST fusion protein of the RhoA binding domain (RBD) of Rhotekin was used. The GST-PBD and GST-RBD beads were provided by Keith Burridge, UNC. Pulled-down proteins from cell lysates (50 µg) were separated using SDS-PAGE and transferred to nitrocellulose. Filters were blocked in 5% nonfat milk in TBS-Tween 20 (0.1%) and incubated overnight with rabbit anti-NrCAM (R&D Systems AF8538, 1:1000), mouse anti-Rac1 (BD Biosciences # 610651, RRID:AB_397978, 1:1000), mouse anti-RhoA (BD Transduction Laboratories #610990, 1:500), or mouse anti-Cdc42 (MAb28-10). In some cases, assays were carried out using kits from Cytoskeleton, Inc. Blots were developed with Advansta enhanced chemiluminescence, and bands were detected within a linear range of film exposure. Bands were quantified by measuring band intensities with ImageJ (NIH) software.

### Synaptoneurosome Preparation and Tiam1 Immunoprecipitation

Synaptoneurosomes were isolated from mouse cortex (P28) as described [28]. Tissue was homogenized in 10 mM HEPES, 1 mM EDTA, 2 mM EGTA, 200 µM Na3VO4, 10 mM NaF, and protease inhibitors. After sonication, the sample was filtered through a 100 μm filter and 5 μm cell strainers, then centrifuged at 1000 × g for 10 min. The pellet was resuspended in 500 μl of RIPA buffer (20 mM Tris pH 7.0, 0.15 M NaCl, 5 mM EDTA, 1 mM EGTA, 1% NP-40, 1% deoxycholate, 0.1% SDS, 200 μM Na3VO4, 10 mM NaF, protease inhibitors), agitated for 40 min, and re-centrifuged at 16,000 × g for 10 min. The supernatant was collected as the synaptoneurosome fraction. Precleared lysates (0.5-1 mg) were incubated with mouse monoclonal anti-TIAM1 (Santa Cruz sc-393315) or normal IgG for 2 hr at 4°C and collected on Protein A/G Sepharose beads. SDS-PAGE and immunoblotting with rabbit NrCAM antibodies was carried out, then filters were stripped and reprobed for Tiam1/2.

### Analysis of Phosphorylated Signaling Intermediates in Spines

For analysis of phosphorylated and non-phosphorylated signaling molecules, mouse cortical neuron cultures expressing EGFP were treated with 5 nM Sema3F-Fc or Fc as described above, fixed in 4% paraformaldehyde (PFA), quenched with 0.1M glycine, permeabilized with 0.1% Triton X-100, and blocked in 10% donkey serum (Sigma #G6767). Cultures were simultaneously stained for EGFP with antibodies directed against GFP and either phospho- or total effector protein. To detect PAK1-3 we used mouse monoclonal α-PAK antibody directed against an epitope mapping between residues 246-470 at the carboxyl terminus (Santa Cruz (A-6) sc-166887). For phosphorylated PAK1-3, staining was carried out using rabbit polyclonal antibody phospho-PAK1 Thr432/ Pak2 Thr402 directed against a synthetic phosphopeptide (Cell Signaling Technology# 2601, RRID:AB_330220). This antibody recognizes phospho-PAK1,2 to a greater extent than PAK3, and does not cross-react with PAK4-6. Phospho-LIMK1/2 was detected with rabbit polyclonal anti-phospho-LIMK1 Thr508/ LIMK2 Thr505 directed against a synthetic phospho-peptide (Thermo Fisher Scientific # PA5-37629, RRID:AB_2554237). For LIMK proteins, we used rabbit polyclonal anti-LIMK antibody directed against LIMK1 residues 600 to the carboxyl terminus (Abcam# ab81046, RRID:AB_2042135). For Cofilin we used rabbit monoclonal anti-Cofilin1/2 antibodies (Cell Signaling Technology# 5175, RRID:AB_10622000), and rabbit monoclonal anti-phospho-Cofilin Ser3 (Cell Signaling Technology# 3313, RRID:AB_2080597). These Cofilin antibodies detect both Cofilin1 and 2, although Cofilin1 is the form that is expressed in neurons. For Myosin II rabbit polyclonal anti-Myosin II light chain (Cell Signaling Technology# 3672, RRID:AB_10692513) and rabbit polyclonal anti-phospho-Ser19 Myosin II light chain (Cell Signaling Technology# 3671, RRID:AB_330248) were used. These antibodies were directed against synthetic peptides common to Myosin light chain II isoforms a,b, and c, and do not recognize cardiac Myosin light chain or Myosin essential light chain. Simultaneous staining for phosphorylated and total effector protein was not possible in most instances due to common host specificity of the antibodies. Secondary Alexa Fluor-conjugated antibodies were obtained from ThermoFisher Scientific, or Jackson Immunochemicals. Imaging was performed using a Zeiss LSM 700 confocal microscope in the UNC Microscopy Services Laboratory as described above.

At least 10 confocal images of apical dendrites from neurons with pyramidal cell morphology in each of 3 or more replicate assays were collected blind to the treatment, as visualized in the EGFP channel (AF488). Identical settings were used to capture images from Fc and Sema3F-Fc treated samples in every case. Deconvolution was carried out using AutoQuant version 3 software (Media Cybernetics) with default settings. To quantify immunofluorescence in spines the EGFP channel was used to create a mask within Bitplane Imaris vXYZ (Oxford Instruments). With the surfaces tool, fluorescence from other channels was set to 0 everywhere outside of the EGFP-containing surface, thus restricting it to the transfected spines being analyzed. Spines were identified as protrusions of at least 0.2 µm on the first branch of apical dendrites. Maximum intensity projections for each channel were generated in FIJI, and total intensity values were gathered for the masked channel using the Time Series Analyzer V3 plugin, which allows placement of multiple regions of interest (ROIs) in an image. A 5×5 pixel square was placed on spine protrusions (mushroom, stubby, thin morphologies) as visualized in the AF488 channel. ROIs were transferred to the masked red channel and the sum of pixel intensity values (the integrated density) within each ROI was measured. These values were then divided by the total number of non-zero pixels within the ROI of the masked image to obtain the average pixel intensity within the spine, which typically occupied less than the total number of pixels in the ROI. Fluorescence intensity values for phosphorylated signaling molecules were divided by the average values for total effector proteins in spines and means (± SEM) were computed for the resulting ratios of phosphorylated/total effector protein. Values for Fc-treated and Sema3F-Fc treated cultures were compared using the nonparametric Mann-Whitney test (2-tailed, p < 0.05). A nonparametric test was used generally because spines responding to Sema3F were not normally distributed. As shown in prefrontal cortex *in vivo* and in cortical neuron cultures, one subpopulation of apical dendritic spines expresses NrCAM and responds to Sema3F, while another subpopulation expresses CHL1 and responds to Sema3B [29, 12]. Two-way ANOVA with Tukey post-hoc testing for multiple comparisons was performed when more than 2 groups were compared such as in all inhibitor experiments.

Spine morphologies were scored as mushroom, stubby, or thin (filopodial) on apical dendrites as defined in [30] and described previously [11]. To compare spine morphologies across different conditions, we applied multinomial logistic regression for computing the log-odds as described [31]. This method was described in detail [11] and briefly, involved a pair of logistic regression models, because the response comprised the three morphological spine types. Because thin spines are the least frequently observed, the models were formulated as log-odds of the outcome to be mushroom rather than thin, and log-odds of the outcome to be stubby rather than thin. The ‘multinom’ function from R ‘nnet’ was employed to estimate the logistic regression models and resulting p-values were reported.

## RESULTS

### Structural Analysis of the NrCAM-Npn2 Interaction in the Sema3F Receptor Complex

A structure-function analysis was performed to investigate how NrCAM engages Npn2 in the Sema3F holoreceptor complex to promote spine pruning. We previously used a molecular modeling approach [12] to predict an interface between the NrCAM Ig1 and Npn2 a1 domains using the ClusPro protein-protein docking server [32,23,22]. We demonstrated that this interface involves an electrostatic interaction between Arg^120^ (R120) in the TAR^120^NER motif of NrCAM Ig1, and Glu^56^ (E56) within Npn2 a1 [12] (Fig. 1 A, B). The model also predicted a closely positioned network of charged residues within NrCAM Ig1 (Asp^82^Asp^84^Lys^85^), termed “DDK network”, stabilized by bonds of length 2.6-2.7 Å at the interface with Npn2 a1. NrCAM Lys^85^ (K85) was predicted to form electrostatic and hydrogen bonds to both NrCAM Asp^82^ (D82) and Asp^84^ (D84) thereby positioning NrCAM Asp^82^ (D82) to interact with Npn2. The acidic Asp^82^ (D82) of NrCAM was predicted to form two additional hydrogen bonds of length 2.7 and 2.8 Å with the basic Arg^31^ (R31) of Npn2, which could provide enhanced stability to the interdomain interaction (Fig. 1B).

To test these predictions, we utilized site-directed mutagenesis to create charge reversal or neutral substitutions designed to disrupt electrostatic interactions or hydrogen bonds. First, Asp^82^ in the NrCAM sequence was mutated to arginine to disrupt intramolecular bonding with Lys^85^ within the DDK network as well as electrostatic interaction with Npn2. WT NrCAM and NrCAM mutant Asp^82^Arg were compared for Npn2 binding by co-immunoprecipitation from lysates of transfected HEK293T cells. NrCAM Asp^82^Arg exhibited reduced Npn2 binding compared to WT NrCAM, as shown by a decrease in the average ratio of Npn2/NrCAM in the immunoprecipitates from 3 replicate experiments normalized to WT Npn2/NrCAM (average ± SEM: 0.170 ± 0.050) (Fig. 1 C). Next, NrCAM Asp^84^ was mutated to alanine to create a neutral substitution that could also impair intramolecular hydrogen bonding within the network, as well as electrostatic interaction with Npn2. This NrCAM mutation also displayed decreased binding to Npn2 compared to WT NrCAM as shown by a decrease in the average ratio of Npn2/NrCAM in the immunoprecipitates from 3 replicate experiments normalized to WT Npn2/NrCAM (average ± SEM: 0.607 ± 0.193) (Fig. 1 C). Finally, NrCAM Lys^85^ was mutated to glutamate to create a charge reversal mutation, which could be the most effective at altering the DDK network by disrupting interactions with both Asp^82^ and Asp^84^. NrCAM Lys^85^Glu exhibited impaired binding to Npn2 as shown by a decrease in the average ratio of Npn2/NrCAM in the immunoprecipitates from 3 replicate experiments normalized to WT Npn2/NrCAM (average ± SEM: 0.181 ± 0.074) (Fig. 1 C). Western blotting of input lysates confirmed that WT and mutant proteins were expressed at approximately equivalent levels in HEK293 cells (Fig. 1 C, lower panel). Correspondingly, a charge reversal substitution in mouse Npn2 from Arg^31^ (R31) to glutamate was designed to test the hypothesis that Arg^31^ is a key residue at the interface, as charge reversal should electrostatically repulse NrCAM Asp^82^ and Asp^84^. Results showed that this Npn2 mutation decreased binding to WT NrCAM, implicating Arg^31^ as a pivotal binding residue in the Npn2 a1 domain as shown by a decrease in the average ratio of Npn2/NrCAM in the immunoprecipitates from 3 replicate experiments normalized to WT Npn2/NrCAM (average ± SEM: 0.496 ± 0.202) (Fig. 1 D). In summary, these results showed that mutating any of the DDK residues compromises NrCAM integrity for Npn2 binding.

One of the most effective NrCAM mutations at perturbing Npn2 binding (NrCAM Lys^85^Glu) was then tested in a functional assay for Sema3F-induced spine retraction on apical dendrites of mouse cortical pyramidal neurons in culture [10, 9]. This established assay measures spine collapse in response to treatment with Sema3F-Fc fusion protein, and correlates well with *in vivo* spine density in showing specificity for NCAM-dependent spine pruning on apical and not basal dendrites [33, 10]. Pyramidal neurons in the cultures are recognized by morphological features including a large cell body with distinguishable apical and basal dendrites. These cells express MATH2 but not GABAergic interneuron markers, and constitute approximately 80% of neuronal cells, similar to the proportion *in vivo*. By DIV13 dendritic spines are evident and form synaptic contacts identified by immunolabeling of SAP102 or PSD95. Neurons cultured from NrCAM null embryos (E14.5) were transfected at DIV11 with vector (pCAGGS-IRES-mEGFP), WT NrCAM, or NrCAM Lys^85^Glu cDNA. Neuronal cultures were treated at DIV14 with Sema3F-Fc or control Fc (5 nM) for 30 min, stained for EGFP, and spine density quantified on apical dendrites from confocal z-stacks after deconvolution. Neurons expressing WT NrCAM exhibited decreased spine density in response to Sema3F-Fc compared to Fc, whereas neurons with vector alone showed no change (Fig. 1 E, F) in accord with prior work [12, 10]. Neurons expressing NrCAM Lys^85^Glu showed no decrease in spine density after Sema3F-Fc addition as determined by 2-way ANOVA in group (WT vs mutant) (p = 0.751) and type (Fc vs. Sema3F-Fc) (p = 0.003). Tukey post-hoc pairwise analysis showed that WT Fc vs. Sema3F-Fc was significantly different (p = 0.021), while mutant Fc vs. Sema3F-Fc was not significant (p = 0.993). Sema3F-Fc treatment was significantly different between WT and mutant cells (p = 0.004WT and mutant NrCAM were appropriately expressed and localized to the neuronal surface, as shown by immunofluorescence staining for NrCAM with antibodies to its extracellular region (red) on unpermeabilized neuronal cells expressing WT and mutant NrCAM (Fig. 1 G).

In summary, these results indicated that the DDK network in NrCAM Ig1 and Arg^31^ in Npn2 a1 are pivotal residues within a charged interface that, together with the NrCAM TARNER motif [10], enable NrCAM to associate with Npn2.

The extended interface between NrCAM Ig1 and Npn2 a1 domains might serve to further stabilize the interaction between Npn2 and PlexA3. To test this possibility an affinity pull-down experiment was designed to assess the ability of NrCAM to promote binding between Npn2 and PlexA3. HEK293T cells were transfected to express PlexA3 and Npn2 in the presence or absence of NrCAM. Sema3F-Fc (5 nM) was added to the cells for 30 min then cells were lysed in Brij98 detergent-containing buffer and immunoprecipitated with Npn2 antibodies. Western blotting for PlexA3 in the Npn2 immune complexes showed that NrCAM increased the levels of associated PlexA3 approximately 4-fold (Fig. 1 H). Input blots verified that equivalent amounts of each protein were expressed in HEK293T cells. These results are consistent with the interpretation that NrCAM increases the stability of the Npn2/PlexA3 interaction.

#### Sema3F Signaling through Rac1 and RhoA GTPases

We have reported that Sema3F induces membrane clustering of the NrCAM/Npn2/PlexA3 complex in cortical neuron cultures within 5-15 min and spine retraction occurs within 15-30 min. We speculated that small GTPases (Rac1, RhoA, Cdc42) might be important effectors, as they are prominent regulators of the actin cytoskeleton. We analyzed Rho-GTPase signaling in mouse cortical neuron cultures following treatment with Sema3F-Fc or Fc proteins. We chose to assay activity at 30 min after treatment since this was when spine pruning was maximal. GTP-bound Rac1, RhoA, and Cdc42 were quantified using GST-pull down assays specific for the activated GTP-bound protein. The relative levels of the GTP-bound protein in the pull-downs were expressed relative to total (GTP- and GDP-bound) Rac1, RhoA, or Cdc42 protein. Sema3F-Fc treatment increased the levels of GTP-bound Rac1 in cultures approximately 3.2-fold over that of Fc-treated control cells in triplicate experiments (Fig. 2 A). Sema3F-Fc increased RhoA-GTP approximately 3.3-fold over Fc controls, while GTP-bound Cdc42 did not change significantly. Pilot experiments showed that Rac1-GTP stimulation occurred early and appeared sustained, increasing approximately 2.5-fold at 2 min, 3.2-fold at 5 min, and 2.6-fold at 10 min. Similarly, RhoA-GTP levels increased and were sustained approximately 2.5-fold at 5 min and 3.8-fold at 10 min. Rac1-GTP and RhoA-GTP levels were relatively unchanged in neuronal cultures from NrCAM null mice comparing the ratios of activities after Sema3F-Fc to Fc treatment (Rac1-GTP, 0.97; and RhoA-GTP, 1.06). This is fully consistent with prior results demonstrating the requirement of NrCAM for Npn2/PlexA3 receptor clustering, Rap1 activation and Sema3F-induced spine retraction [10, 12].

**Fig. 2.**
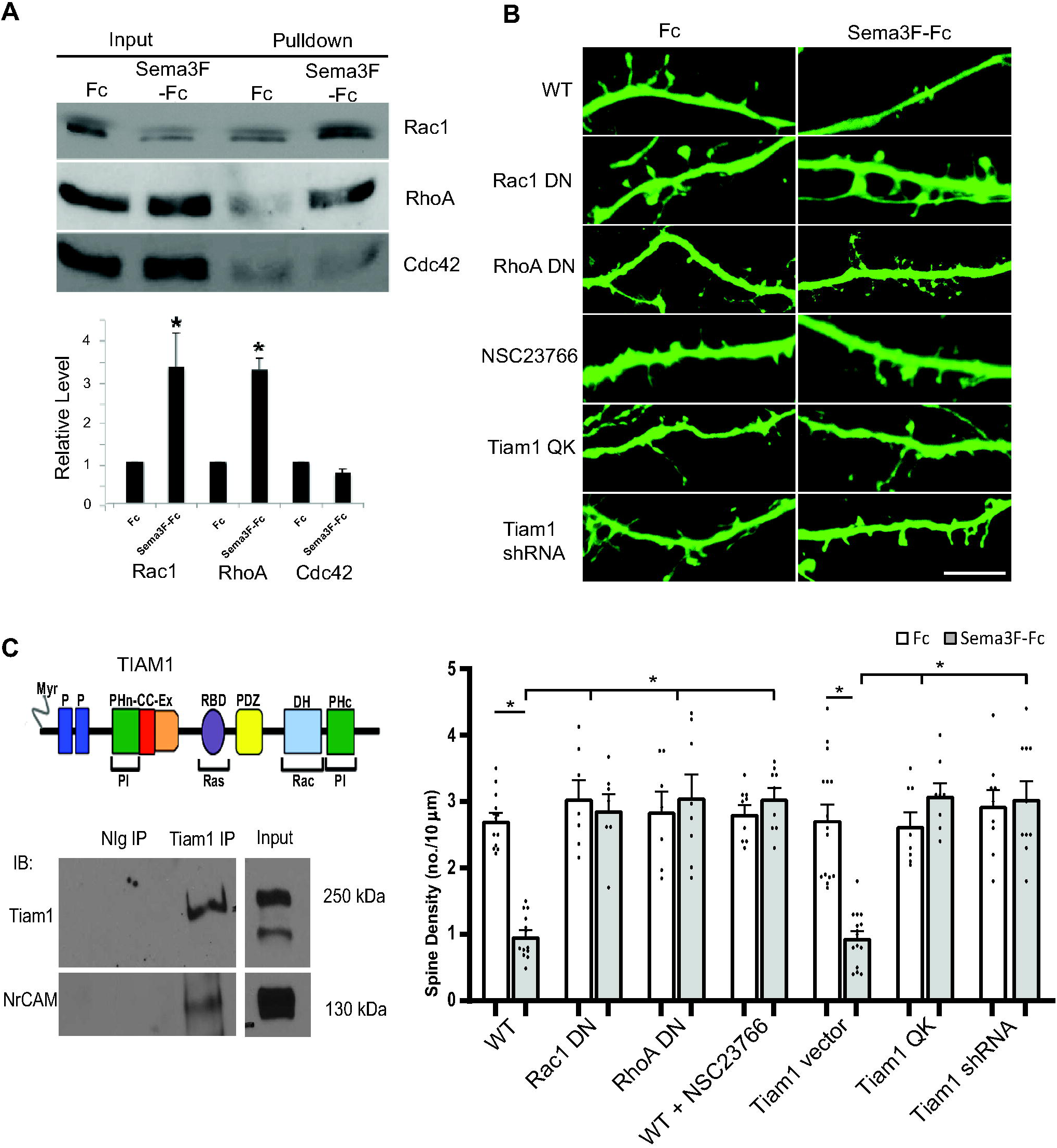
Sema3F-induced spine remodeling in cortical neuron cultures is mediated through Rac1 and RhoA GTPase. A. Rac1, RhoA, and Cdc42 pulldown assays from cortical neuron cultures (DIV14) showed that Sema3F-Fc induced GTP-loading/activation of Rac1 and RhoA but had no effect on Cdc42 after 30 min. GTP-bound GTPases were pulled down from equal amounts of lysate protein from Sema3F-Fc or Fc-treated cells with GST-fusion effector proteins on Sepharose beads. Pull-downs and input cell lysates were analyzed on Western blots using antibodies to total Rac1, RhoA, or Cdc42 protein. A representative blot from 3 experiments is shown. Densitometric band intensities of pull-downs from 3 replicate assays were divided by the amount of input protein, and relative mean levels plotted (± SEM). P-values were for Rac1-GTP (*p = 0.005), RhoA-GTP (*p = 0.001), Cdc42-GTP (p = 0.060) (t-test, 2-tailed). B. WT cortical neurons were transfected with pCAGG-IRES-EGFP treated with Fc or Sema3F-Fc for 30 min at DIV14, immunostained for EGFP, and apical dendrites imaged confocally. Cultures were co-transfected with dominant negative (DN) mutant Rac1 or RhoA plasmids or treated with NSC23766 to inhibit Tiam1 activation of Rac1. Alternatively, cells were transfected with plasmids expressing Tiam1QK or Tiam1 shRNA. Scale bar = 5µm. Below, quantification of mean spine density ± SEM on apical dendrites (n > 300 spines and ≥ 10 neurons per condition). Each point represents the mean spine density per neuron. 2-way ANOVA with Tukey post-hoc pairwise comparisons showed statistical significance (WT Fc vs. Sema3F-Fc, p = 0.001; WT Sema3F-Fc vs. Rac1 DN + eGFP Fc, Rac1 DN + eGFP Sema3F-Fc, RhoA DN + eGFP Fc, RhoA DN + eGFP Sema3F-Fc, WT + NSC23766 Fc, or WT + NSC23766 Sema3F-Fc, p = 0.001; and Tiam1 vector + eGFP Fc vs. Sema3F-Fc, p = 0.001; Tiam1 vector + eGFP Sema3F-Fc vs. Tiam1 QK + eGFP Fc, p = 0.020; Tiam1 vector + eGFP Sema3F-Fc vs. Tiam1 QK + eGFP Sema3F-Fc, Tiam1 shRNA + eGFP Fc, or Tiam1 shRNA + eGFP Sema3F-Fc, p = 0.001). C. Schematic of TIAM1 domain structure showing myristylated N-terminus (Myr), binding sites for phosphoinositides (PI) (PHn-CC-Ex and PHc), Ras (RBD), PDZ, and Rac1 (dbl homology, DH). Co-immunoprecipitation of Tiam1 and NrCAM from postnatal mouse synaptoneurosomes, shown by immunoprecipitation (IP) with normal Ig (NIg) or Tiam1 antibodies, followed by immunoblotting (IB) for Tiam1 and reprobing blots for NrCAM. Input lysate is shown at right. The blots shown are representative of 3 experiments (Supplementary Figure 2).

To further probe a functional role for Rac1 and RhoA in Sema3F-induced spine density regulation, we analyzed the effect of expressing dominant negative (DN) mutants and pharmacological inhibitors in cortical neuron cultures expressing pCAGG-IRES-EGFP. RhoA family GTPases are activated by GTP loading catalyzed by guanine nucleotide exchange factors (GEFs). The dominant negative (DN) Rac1 mutant Rac1-17N contains a threonine to asparagine substitution at position 17, which sequesters Rac-GEFs, thus preventing Rac1 from becoming activated. DN Rac1-17N effectively inhibited Sema3F-induced spine retraction but did not significantly alter spine density in Fc-treated cells (Fig. 2 B). Similarly, expression of DN RhoA-G17A, which prevents RhoA activation by sequestering Rho-GEFs, blocked Sema3F-Fc-induced spine retraction in neuronal cultures (Fig. 2 B).

Although Rac1 can be activated by multiple GEFS, we focused on Tiam1/2, which activates Rac1/2 selectively and localizes to spines [24]. Tiam1/2 is a multi-domain GEF that binds Rac1, phosphoinositides, and Ras (Fig. 2 C). Cortical neuron cultures were pretreated with NSC23766 (50 µM), a selective inhibitor of Rac1 activation by Tiam1/2-GEF or the related GEF Trio [34], and assayed for spine retraction by Sema3F-Fc. NSC23766 effectively impaired Sema3F-induced spine loss (Fig. 2 B). To specifically address the role of Tiam1, neurons were transfected with plasmids expressing the Tiam1 GEF-dead mutant Q^1191^K^1195->^AA (Tiam1 QK) or Tiam1 shRNA, which downregulates Tiam1 but not Tiam2. Both Tiam1 QK and Tiam1 shRNA inhibited Sema3F-Fc induced spine retraction compared to empty vector (Fig. 2 B). We next examined the association of Tiam1 with NrCAM in synaptoneurosomes, a subcellular fraction of mouse cortex (P28) that is enriched in pre- and post-synaptic terminals [28]. Tiam1 was immunoprecipitated from synaptoneurosome lysates and found to co-immunoprecipitate with NrCAM (Fig. 2 C and Supplemental Figure 2). However, Tiam1 and NrCAM did not co-immunoprecipitate from transfected HEK293 cells (not shown), suggesting that the interaction may be indirect. The Tiam1 PHn-CC-Ex domain can bind the actin adaptor Ankyrin [35], which reversibly engages the cytoplasmic tail of NrCAM [36]. Neither Ankyrin-B nor Ankyrin-G co-immunoprecipitated with NrCAM and Tiam1 from synaptoneurosomes despite their presence in the fraction (not shown). These results demonstrate a role for Rac1 and RhoA GTPases in the molecular mechanism of Sema3F-induced spine remodeling.

#### Sema3F Downstream Signaling through Rac1, PAK1-3, LIMK1/2, and Cofilin1

Principal downstream effectors of Rac1 include the p21-activated kinases (PAKs), a family of serine/threonine kinases that regulate actin remodeling [37]. Group I PAKs (PAK1-3) are closely related in structure, whereas Group II PAKs 4-6 are more divergent. PAK1-3 is expressed in neurons where it regulates excitatory synaptic functions including spine morphology and learning/memory [38]. Autophosphorylation at Thr423(PAK1), Thr402(PAK2), or Thr421(PAK3) is a hallmark of PAK1-3 activation in dendritic spines [39]. Because spines compartmentalize small GTPases for synaptic transmission and plasticity [40], we examined the activation of signaling intermediates in spines by immunofluorescence. To determine if PAK1-3 becomes activated within apical dendritic spines upon Sema3F stimulation, cortical neuron cultures were transfected with pCAGG-IRES-EGFP then treated with Sema3F-Fc or Fc for 10 minutes. Neurons were processed for immunofluorescence staining with antibodies directed against phospho-PAK1 Thr432, which recognizes all 3 group I isoforms, and antibodies to PAK1-3 protein. Phospho-PAK1-3 and PAK1-3 immunolabeling was evident along apical dendrites as discrete puncta (Fig. 3 A), which at higher magnification (lower panels) were morphologically identifiable as spines. When images were captured at the same settings, phospho-PAK immunofluorescence was seen to increase following Sema3F-Fc treatment compared to Fc (Fig. 3 A). Phospho- and PAK1-3 protein immunofluorescence was quantified in individual spines by restricting pixel density measurements to masked GFP-labeled spines after deconvolution of confocal z-stacks. Relative levels of phospho-PAK1-3 to PAK1-3 protein were significantly increased by Sema3F-Fc compared to Fc at 10 min (p = 0.011, Mann-Whitney 2-tailed test) (Fig. 3 B). Average PAK1-3 protein levels after Sema3F-Fc treatment were somewhat decreased compared to Fc (mean pixel intensity ± SEM; Sema3F-Fc 124.9 ± 8.9, Fc 172.0 ± 11.7, p = 0.002, Mann-Whitney 2-tailed test) but this change was offset by a greater increase in phospho-PAK levels (mean pixel intensity ± SEM; Sema3F-Fc 266.5 ± 19.1, Fc 194.1 ± 14.0, p = 0.001, Mann-Whitney 2-tailed test). The protein change might be due to degradation or translocation. PAK1-3 phosphorylation in spines, and that of other downstream effectors, could not be verified by Western blotting due to the presence of many nonneuronal cells in the culture.

**Fig. 3.**
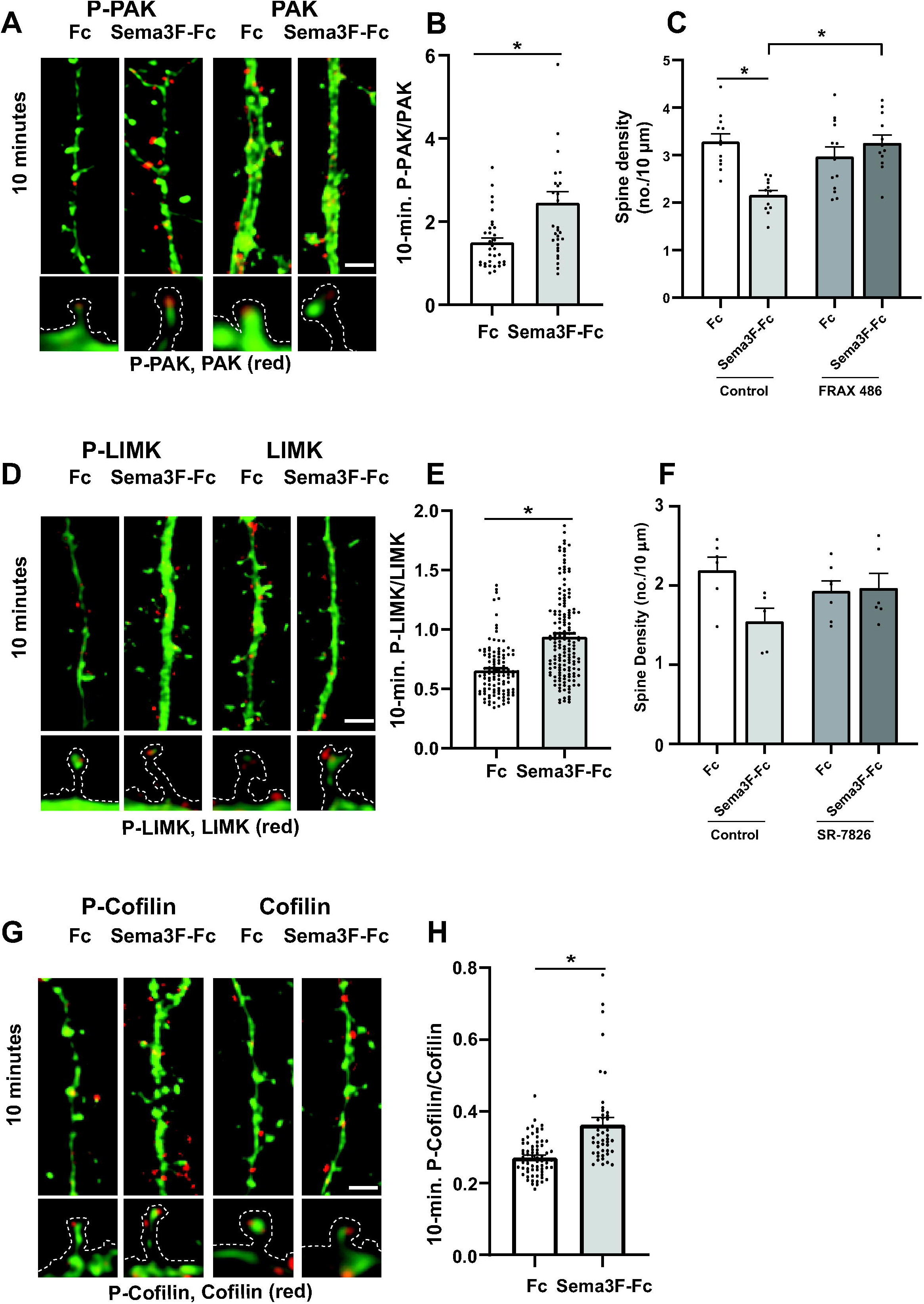
Sema3F induces spine retraction in cortical neuron cultures through PAK, LIMK, and Cofilin phosphorylation. A. Representative images of EGFP-labeled apical dendrites with spines immunostained for phospho-PAK1-3 (red) or PAK1-3 protein (red) after 10 min of Sema3F-Fc or Fc treatment. Scale bar = 2µm. Optically zoomed images of spines outlined are shown below. B) Quantification of pixel intensity of phospho-PAK1-3 in spines divided by PAK1-3 protein intensity at 10 minutes of treatment with Sema3F-Fc or Fc. Relative levels of phospho-PAK1-3 to PAK1-3 protein were significantly increased by Sema3F-Fc compared to Fc at 10 min (p = 0.011, Mann-Whitney 2-tailed test). Each point represents phospho-PAK/PAK per spine from n > 100 spines and ≥ 6 neurons per condition. C) Quantification of spine density on apical dendrites in neuronal cultures pretreated for 30 minutes with control vehicle (0.01% DMSO) or PAK inhibitor FRAX 486. FRAX 486 (500 nM) inhibited Sema3F-Fc induced spine retraction as determined by 2-way ANOVA with Tukey post-hoc pairwise comparisons (Control Fc vs. Sema3F-Fc, p = 0.001; Control Sema3F-Fc vs. FRAX 486 Sema3F-Fc, p = 0.001; FRAX 486 Fc vs. FRAX 486 Sema3F-Fc, p = 0.611). Each point represents the mean spine density per image (n > 300 spines and ≥ 10 neurons per condition). D) Representative images of eGFP-labeled apical dendrites and spines immunostained for phospho-LIMK1/2 (red) or LIMK1/2 protein (red) after 10 minutes of Sema3F-Fc or Fc treatment. Scale bar = 2µm. Optically zoomed images of spines outlined are shown below. E) Quantification of pixel intensity of phospho-LIMK1/2 in spines divided by LIMK1/2 protein intensity at 10 minutes of treatment with Sema3F-Fc or Fc. A significant increase in mean phospho-LIMK/LIMK was observed after 10 minutes of Sema3F-Fc treatment compared to Fc (*p = 0.0001, Mann-Whitney 2-tailed test). Each point represents the p-LIMK/LIMK ratio per spine (n> 250 spines and n ≥ 10 neurons per condition. F) Quantification of spine density on apical dendrites in neuronal cultures pretreated for 30 minutes with vehicle control (0.01% DMSO) or with LIMK1/2 inhibitor SR-7826. SR-7826 (1 µM) decreased Sema3F-Fc reduced spine retraction compared to Fc control although not reaching statistical significance by 2-way ANOVA with Tukey post-hoc pairwise comparisons (Control Fc vs. Sema3F-Fc, p = 0.063; SR-7826 Fc vs. SR-7826 Sema3F-Fc, p = 0.998). Each point represents the mean spine density per neuron (n > 300 spines and ≥ 6 neurons per condition). G) Representative images of apical dendrites with spines of EGFP-labeled cortical neurons showing immunostaining of phospho-Cofilin1 (red) or Cofilin1 (red) after 10 minutes of Sema3F-Fc or Fc treatment. Scale bar = 2µm. Optically zoomed images of spines outlined are shown below. H) Quantification of pixel intensity of phospho-Cofilin1/2 in spine heads divided by Cofilin1 protein intensity at 10 minutes of treatment with Sema3F-Fc or Fc. Mean p-Cofilin1/Cofilin1 was significantly increased after Sema3F-Fc treatment (*p = 0.0001, Mann-Whitney 2-tailed test). Each point represents phospho-Cofilin1/Cofilin1 per spine (n > 300 spines and 6 neurons per condition).

To assess whether PAK1-3 kinase activity was required for Sema3F-induced spine retraction, neuronal cultures were pretreated with FRAX 486, a small molecule pyrido-pyrimidinone inhibitor of PAK1-3 kinase that shows selectivity over Group 2 PAKs [26]. FRAX 486 competes at the ATP binding site of PAK1-3 and inhibits purified PAK kinase with an IC50 of 14-39 nM [41]. FRAX 486 (500 nM) blocked Sema3F-Fc induced spine retraction as determined by 2-way ANOVA in group (Control vs. FRAX 486) (p = 0.025), in type (Fc vs. Sema3F-Fc) (p = 0.018), and group:type interaction (p = 0.001). Tukey post-hoc pairwise analysis showed that the difference in spine density for Control Fc vs. Sema3F-Fc was significant (p = 0.001), while that of FRAX 486 Fc vs. Sema3F-Fc was not significant (p = 0.611) (Fig. 3 C). In addition, Sema3F-Fc treated Control vs FRAX 486 cells showed a significant difference in spine density (p = 0.001). Supplementary Figure 1 shows scatter plots for the effect of the inhibitor on spine density.

Dendritic spines have different morphologies classified as mushroom, stubby, and thin or filopodial based on the relative size of the spine head and neck (Peters and Kaiserman-Abramof, 1970). These types are dynamically interchangeable from immature thin spines to mature mushroom spines with mature synapses. We quantified the percentages of spines with mushroom, stubby, and thin morphology on apical dendrites of EGF-expressing neuronal cells (Fc-treated controls) in the cultures after inhibitor treatment. There was no significant difference for spines to have mushroom morphology instead of thin or stubby in neurons treated with FRAX 486 versus untreated neurons, as demonstrated by multinomial logistic regression analysis (Supplementary Table 1).

PAK1-3 kinases can regulate actin cytoskeletal remodeling by phosphorylating and activating the serine/threonine kinase LIM kinase (LIMK1/2) on Thr508, which in turn phosphorylates and inactivates the actin severing protein Cofilin 1/2 on Ser3 [42]. Cofilin-mediated severing modulates actin dynamics in a complex way by causing F-actin disassembly as well as providing new ends for F-actin polymerization [43, 44]. LIMK1 is expressed primarily in the central nervous system, whereas LIMK2 is ubiquitous [43].

To determine if LIMK1/2 is phosphorylated and activated within dendritic spines upon Sema3F stimulation, cortical neuron cultures were treated with Sema3F-Fc or Fc for 10 min, a time when PAK1-3 phosphorylation was first observed, then processed for immunofluorescence staining with antibodies against phospho-Thr508 LIMK, which recognizes LIMK1/2, and antibodies recognizing LIMK protein. Phospho-LIMK1/2 and LIMK labeling on apical dendrites was seen within discrete spine puncta, and phospho-LIMK1/2 appeared elevated after 10 min of Sema3F-Fc addition, correlating with PAK phosphorylation/activation (Fig. 3 D). The ratio of phospho-LIMK1/2 to LIMK in spines significantly increased in Sema3F-Fc treated cells compared to Fc controls (Mann-Whitney 2-tailed test, p = 0.0001) (Fig. 3 E). No significant changes were seen in LIMK protein levels after Sema3F-Fc treatment compared to Fc (mean pixel intensity ± SEM; Sema3F-Fc 93.0 ± 3.1, Fc 91.2 ± 2.5, p = 0.678, Mann-Whitney 2-tailed test).

To ask whether LIMK activity was required for Sema3F-induced spine retraction, neuronal cultures were pretreated with SR-7826, a bis-aryl urea compound that is a selective LIMK1/2 inhibitor [45]. SR-7826 is a high potency inhibitor (43 nM IC50 for LIMK1) and is more than 100-fold selective over ROCK1/2 [46]. SR-7826 (1 µM) reduced Sema3F-Fc induced spine retraction compared to Fc controls as supported by 2-way ANOVA in group (Control vs. SR-7826) (p = 0.769), in type (Fc vs. Sema3F-Fc) (p = 0.106), and group:type interaction (p = 0.049). Although Sema3F-Fc decreased spine density in Control neurons (fig. 3F), Tukey post-hoc pairwise analysis indicated that Control Fc vs. Sema3F-Fc did not reach statistical significance (p = 0.063). However, the 95% confidence interval (−1.315 to 0.026) was just barely exceeded (did not include zero). In SR-7826 treated neurons the difference between Fc vs. Sema3F-Fc was insignificant to a much greater degree (p = 0.998) (Fig. 3 F). Supplementary Figure 1 shows scatter plots for effect of the inhibitor on spine density. Spine morphology was not altered by the inhibitor (Supplementary Table 1).

Cofilin1 is expressed in neurons and other cell types, whereas Cofilin2 is restricted to muscle [47]. To determine if Cofilin1 in cortical neurons was phosphorylated and activated within dendritic spines upon Sema3F stimulation, cultures were treated with Sema3F-Fc or Fc and processed for immunofluorescence labeling with antibodies directed against phospho-Ser3 Cofilin1/2 or antibodies against Cofilin1/2 protein for 10 min. Phospho-Cofilin1 and Cofilin1 protein on apical dendrites was evident in spine puncta, and phospho-Cofilin1 appeared to increase after Sema3F-Fc treatment (Fig. 3 G). Quantification revealed that the ratio of phospho-Cofilin1 to Cofilin1 in spines increased to a small but significant extent by 10 min of Sema3F-Fc addition (Mann-Whitney 2-tailed test, p = 0.0001) (Fig. 3 H). No significant changes in Cofilin1/2 levels after Sema3F-Fc treatment compared to Fc were seen (mean pixel intensity ± SEM; Sema3F-Fc 84.8 ± 7.0, Fc 109.5 ± 13.0, p = 0.101, Mann-Whitney 2-tailed test).

In summary, the results are consistent with the interpretation that Sema3F signaling in cortical neuron cultures sets in motion a pathway downstream of Rac1 involving phosphorylation of PAK1-3, LIMK1/2, and Cofilin1 leading to spine retraction.

#### Myosin II and ROCK1/2 Mediate Sema3F-induced Spine Retraction

Although RhoA has multiple downstream targets, the Ser/Thr kinase ROCK1/2 is a potential effector for Sema3F-induced spine retraction, because it is an integration point for signaling pathways that regulate actomyosin contraction [48]. Non-muscle MLC II, a principal target of ROCK1/2, crosslinks actin filaments and induces ATPase-dependent contraction when phosphorylated on Ser19 to generate tension in actin cortical networks [49]. ROCK1/2 also phosphorylates and inactivates MLC phosphatase, further increasing the levels of MLC II phosphorylation [48]. To investigate whether Myosin II becomes phosphorylated in response to Sema3F, MLC II phosphorylation on Ser19 was assayed by immunostaining neuronal cultures treated with Sema3F-Fc or Fc with phospho-specific MLC II antibodies. Phospho-MLC II and MLC II labeling on apical dendrites was generally localized to discrete spine puncta, where phospho-MLC II increased 10 min after Sema3F-Fc addition (Fig. 4 A). Quantification showed the ratio of phospho-MLC II to MLC II in spines was not affected 2 min after Sema3F-Fc addition (p = 0.547, Mann-Whitney 2-tailed test) but increased to a small but significant extent after 10 min (p = 0.031) (Fig. 4 B). No significant changes in MLC II levels after Sema3F-Fc treatment compared to Fc were seen at 2 minutes (mean pixel intensity ± SEM; Sema3F-Fc 108.8 ± 2.9, Fc 108.1 ± 2.8, p = 0.802, Mann-Whitney 2-tailed test) or 10 minutes (mean pixel intensity ± SEM; Sema3F-Fc 49.6 ± 1.4, Fc 48.2 ± 1.4, p = 0.329, Mann-Whitney 2-tailed test).

**Fig. 4.**
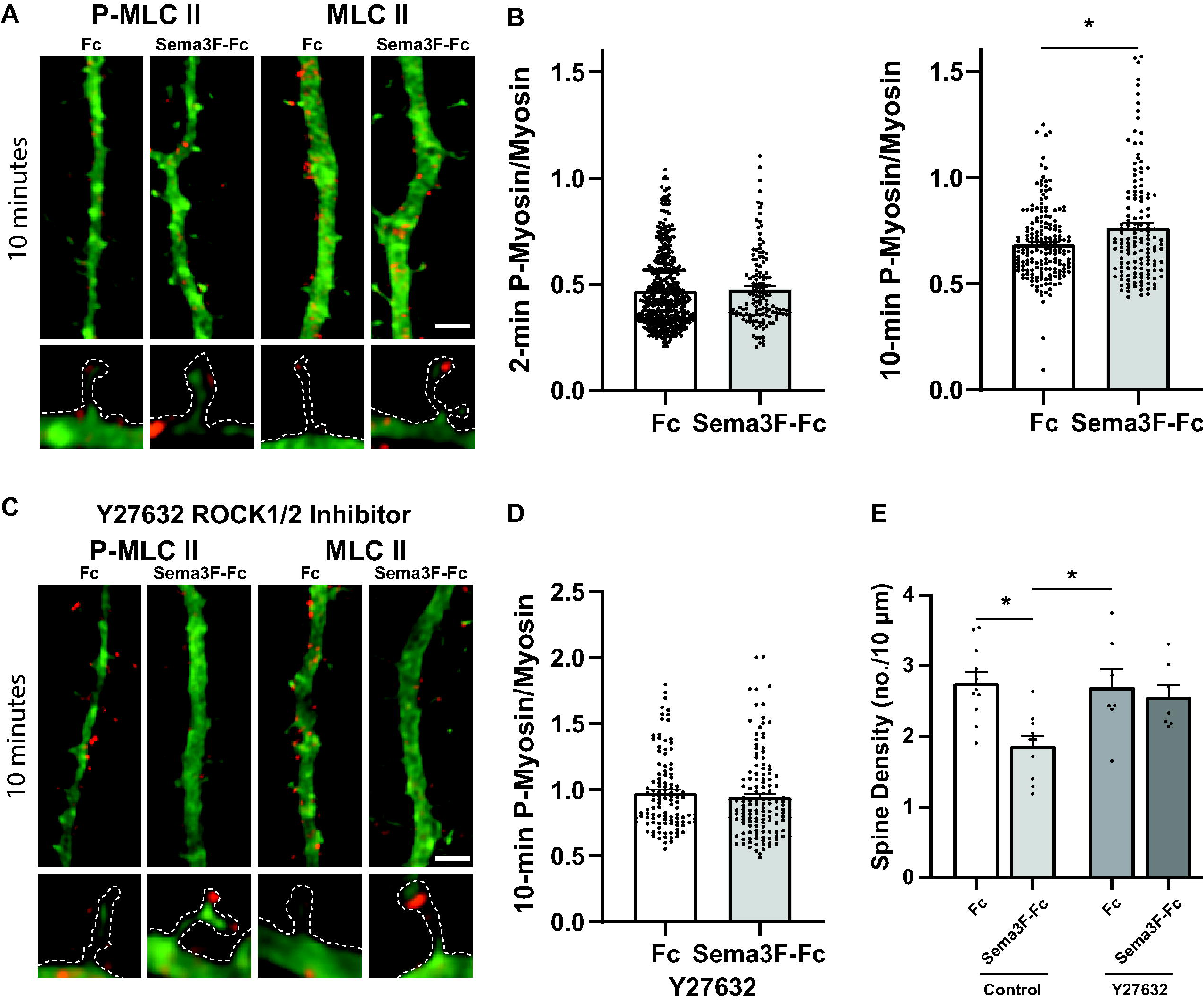
Sema3F induces spine retraction in cortical neurons through Myosin II and ROCK1/2. A) Representative images of EGFP-labeled cortical neurons immunostained for phospho-MLC II (red) or MLC II protein (red) after 10 minutes of Sema3F-Fc or Fc treatment. Scale bar = 2µm. Optically zoomed images of spines outlined are shown below. B) Quantification of pixel intensity of phospho-MLC II in spines divided by MLC II protein intensity at 2 and 10 minutes of treatment with Sema3F-Fc or Fc. A significant increase in mean phospho-MLC II/MLC II is seen at 10 min after Sema3F-Fc addition. P-values (Mann-Whitney 2-tailed test) were for 2-minute phospho-MLC II/MLC II (p = 0.547) and 10-minute phospho-MLC II/MLC II (*p = 0.031). Each point represents the phospho-MLC II/MLC II ratio per spine (n > 270 spines and ≥ 10 neurons per condition). C) Pretreatment of cortical neurons with ROCK1/2 inhibitor Y27632 (10µM) for 1 hour inhibits Sema3F-induced MLC II phosphorylation. Representative images of EGFP-labeled cortical neurons showing apical dendrites with spines immunostained for phospho-MLC II (red) or MLC II protein II (red) after 10 min of Seam3F-Fc or Fc treatment. Optically zoomed images of spines outlined are shown below. D) Quantification of pixel intensity of phospho-MLC II in spines divided by MLC II protein intensity in Y27632-treated neurons at 10 minutes of treatment with Sema3F-Fc or Fc. Phospho-MLC II/MLC II was unaltered in Y27632-treated neurons after Sema3F-Fc treatment (p = 0.234, Mann-Whitney 2-tailed test). Each point represents the phospho-MLC II/MLC II ratio per spine from n > 240 spines and ≥ 10 neurons per condition. E) Quantification of spine density in neurons pretreated for 1 hour with Y27632 (10µM) before 30 min of Sema3F-Fc or Fc treatment. Y-27632 inhibited Sema3F-Fc induced spine retraction compared to Fc as determined by 2-way ANOVA with Tukey post-hoc pairwise comparisons (Control Fc vs. Sema3F-Fc, p = 0.003; Control Sema3F-Fc vs. Y27632 Sema3F-Fc, p = 0.018; Y27632 Fc vs. Y27632 Sema3F-Fc, p = 0.965). Each point represents the mean spine density per neuron (n > 350 spines and ≥ 12 neurons per condition).

The RhoA substrate ROCK 2 is preferentially expressed in brain, whereas ROCK 1 is present in many tissues [48]. To ask whether ROCK1/2 mediates Sema3F-induced Myosin II phosphorylation, neuronal cultures were treated with Y-27632, a selective cyclohexane carboxamide inhibitor of ROCK1/2 [50]. Y-27632 efficiently inhibits purified ROCK1 (Ki = 220 nM) and ROCK2 (Ki = 300 nM) by competing with ATP binding at the active site. Y-27632 (10 µM) prevented the increase in phospho-Myosin II/Myosin II by Sema3F-Fc in spines at 10 min (p = 0.534, Mann-Whitney 2-tailed test) compared to Fc controls (Fig. 4 C, D). No significant changes in Myosin II protein levels after Sema3F-Fc treatment was seem compared to Fc (mean pixel intensity ± SEM; Sema3F-Fc 46.7 ± 1.5, Fc 42.0 ± 1.2, p = 0.061, Mann-Whitney 2-tailed test). To evaluate the role of ROCK1/2 in Sema3F-induced spine retraction, neuronal cultures were pretreated with Y-27632 (10 µM) and assayed for Sema3F-Fc induced spine retraction at 30 min. Y-27632 effectively blocked Sema3F-Fc induced spine retraction compared to Fc as determined by 2-way ANOVA in group (Control vs. Y-27632) (p = 0.120), in type (Fc vs. Sema3F-Fc) (p = 0.003), and group:type interaction (p = 0.047). Tukey post-hoc pairwise analysis showed that Control Fc vs. Sema3F-Fc was significantly different (p = 0.003), while Y-27632 Fc vs. Sema3F-Fc was not significant (p = 0.965) (Figure 4 E). Supplementary Figure 1 shows scatter plots for effect of the inhibitor on spine density. Spine morphology was unaffected by the inhibitor (Supplementary Table 1) in agreement with other studies [51]. Blebbistatin (25-50 µM, 30-60 min) was not used to assess the role of Myosin II ATPase, because it caused spines in the cortical neuron cultures to adopt a thin, filopodial morphology, as also noted in hippocampal cultures [52].

In summary these results suggest that Sema3F triggers downstream signaling through RhoA and ROCK-mediated phosphorylation of Myosin II necessary to achieve spine retraction.

#### Evidence for Actin Remodeling in Sema3F-induced Spine Retraction

Results presented above implicate signaling pathways that promote both actin assembly (Rac1-PAK-LIMK-Cofilin) and actomyosin contraction (RhoA-ROCK-Myosin II) in Sema3F-induced spine pruning. These opposing forces may coordinate to generate cytoskeletal tension leading to actin remodeling and spine loss as suggested for nonneuronal cells [53,54,49]. To explore a role for actin remodeling in Sema3F-induced spine retraction, we assayed the effect of Latrunculin A (LatA), a cyclic compound that inhibits actin filament formation by binding G-actin [55]. Pretreatment of cortical neuron cultures for 20 min with 2 µM LatA significantly reduced Sema3F-induced spine loss compared to Fc as determined by 2-way ANOVA in group (Control vs. LatA) (p = 0.001), in type (Fc vs. Sema3F-Fc) (p = 0.087), and group:type interaction (p = 0.038). Tukey post-hoc pairwise analysis showed that Control Fc vs. Sema3F-Fc was significantly different (p = 0.042), while LatA Fc vs. Sema3F-Fc was not significant (p = 0.992) (Fig. 5 A, C). Neuronal cultures were next pretreated for 20 min with Jasplakinolide, a membrane-permeable cyclic peptide that inhibits actin depolymerization [55]. Jasplakinolide (JSK; 1 µM) significantly reduced Sema3F-induced spine loss as determined by 2-way ANOVA in group (Control vs. JSK treated) (p = 0.668), in type (Fc vs. Sema3F-Fc) (p = 0.037), and group:type interaction (p = 0.005). Tukey post-hoc pairwise analysis showed that Control Fc vs. Sema3F-Fc was significantly different (p = 0.004), while JSK Fc vs. Sema3F-Fc was not significant (p = 0.999) (Fig. 6 B, D). Supplementary Figure 1 shows a scatter plot for effect of the inhibitor on spine density. Neither LatA nor JSK altered spine morphology at the concentrations used in the cortical neuron cultures in accord with lack of morphology change in hippocampal neuron cultures (Supplementary Table 1) [56].

**Fig. 5.**
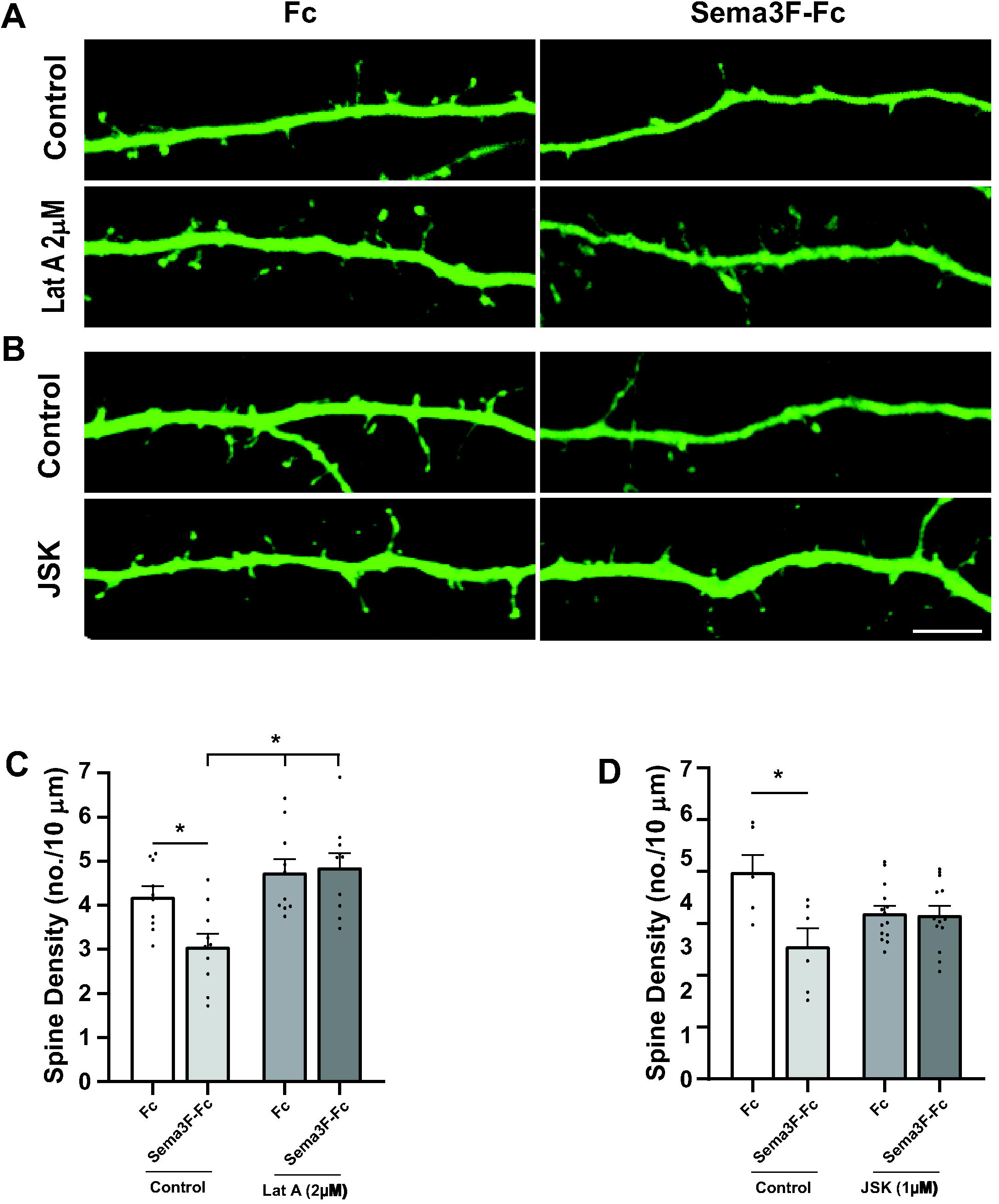
Sema3F-induced spine retraction requires both polymerization and de-polymerization of F-actin. A) Cortical neuronal cultures were pretreated for 20 minutes with an inhibitor of actin polymerization Latrunculin A (Lat A) prior to treatment with Fc or Sema3F-Fc for 30 minutes. Representative images show Sema3F-Fc decreased spine density on apical dendrites of vehicle control cells (0.01% DMSO) but not Lat A (2 µM) treated cells. Scale bar = 5µm. B) Cortical neuronal cultures were pretreated for 20 minutes with an inhibitor of actin depolymerization Jasplakinolide (JSK; 1 µM) prior to Fc or Sema3F-Fc treated for 30 minutes. Representative images show Sema3F-Fc decreased spine density on apical dendrites of vehicle treated control cells (0.01% DMSO) but not JSK (1 µM) treated cells. Scale bar = 5µm. C) Quantification of spine density (mean ± SEM) in cortical neurons treated with vehicle control or Lat A (2 µM). LatA significantly reduced Sema3F-induced spine loss compared to Fc as determined by 2-way ANOVA with Tukey post-hoc pairwise comparisons (Control Fc vs. Sema3F-Fc, p = 0.042; Control Sema3F-Fc vs. LatA Fc, p = 0.001; Control Sema3F-Fc vs. LatA Sema3F-Fc, p = 0.001; LatA Fc vs. LatA Sema3F-Fc, p = 0.992). Each point represents the mean spine density per neuron (n > 300 spines and ≥ 12 neurons per condition). D) Quantification of spine density (mean ± SEM) in cortical neurons treated with vehicle control or JSK (1µM). JSK significantly reduced Sema3F-induced spine loss compared to Fc as determined by 2-way ANOVA with Tukey post-hoc pairwise comparisons (Control Fc vs. Sema3F-Fc, p = 0.003; JSK Fc vs. JSK Sema3F-Fc, p = 0.992). Each point represents the mean spine density per neuron (n > 1000 spine and ≥ 20 neurons per condition).

**Fig. 6.**
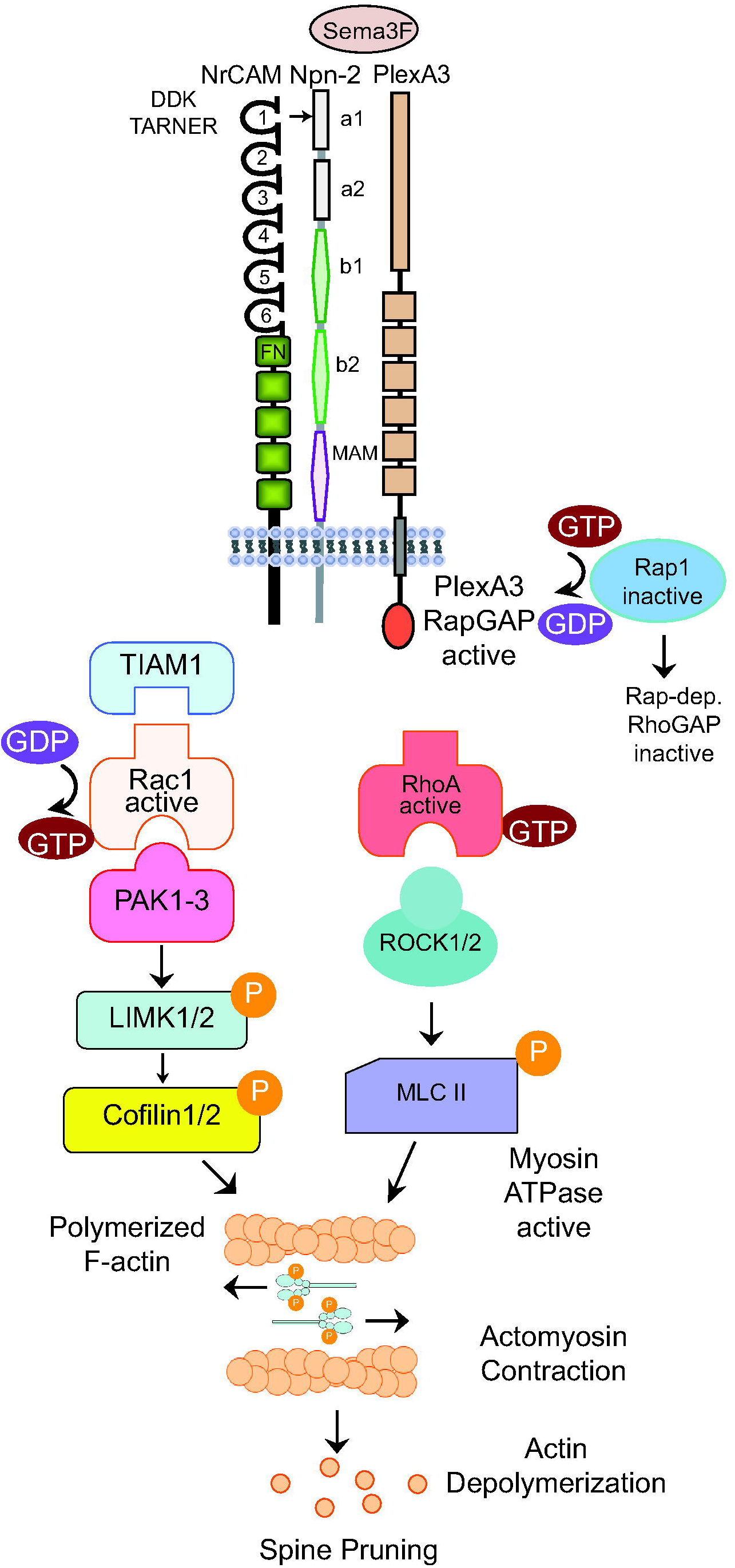
Postulated Dual Signaling Pathway for Sema3F-induced Spine Pruning through Rac1 and RhoA. Sema3F induced signaling begins with binding between the NrCAM-Npn2-PlexA3 holoreceptor and ligand Sema3F. Sema3F induces clustering and conformational changes that initiates the RapGAP activity of the intracellular portion of PlexA3, leading to the inactivation of Rap1. It is speculated that a Rap-dependent RhoGAP is inactivated allowing RhoA-GTP to be active. GTP-bound RhoA binds ROCK1/2 which then activates Myosin II by phosphorylating the myosin light chain (MLC). Phosphorylated MLC II induces actomyosin contraction which may create tension that leads to actin depolymerization. To provide the framework necessary for contractile force, Tiam1 is recruited to exchange GDP for GTP on Rac1. Rac1 then activates PAK1-3. PAK can then phosphorylate LIMK1/2, which phosphorylates Cofilin1, inactivating it. Cofilin inactivation leads to elongation of actin filaments providing the framework for tension generation.

Sema3F-Fc induces co-clustering of Npn2 and PlexA3 on apical dendrites as shown in cortical neuron cultures by immunofluorescence co-localization [12]. Pretreatment of cortical neuron cultures with Lat A (2 µM) had no effect on Sema3F-induced co-clustering of Npn2 and PlexA3 in this assay (not shown), suggesting that actin polymerization is not a factor in receptor clustering. These findings suggested that Sema3F signaling culminates in actin cytoskeletal remodeling necessary for spine retraction.

## DISCUSSION

The structure-function studies presented here demonstrate that NrCAM engages Npn2 within the Sema3F receptor complex via a network of charged residues in the NrCAM Ig1 and Npn2 a1 domains. This extended interface provides stability to the interaction between Npn2 and PlexA3 essential for spine pruning. We also show that Sema3F sets in motion dual Rac1 and RhoA GTPase signaling pathways that converge at the level of the actin cytoskeleton to promote spine elimination.

A molecular modeling and mutagenesis approach revealed that NrCAM Ig1 and Npn2 a1 domains are principal determinants of constitutive binding necessary for spine pruning [12]. In the present studies we identified a closely positioned network of charged residues Asp^82^ Asp^84^ Lys^85^ (DDK) in NrCAM Ig1, and Arg^31^ in Npn2 a1, which are required for Npn2 binding and Sema3F-induced spine retraction. Modeling predicted that NrCAM Lys^85^ hydrogen bonds intramolecularly with NrCAM Asp^82^ and Asp^84^, positioning these residues for electrostatic interaction with Npn2 Arg^31^. Substitution mutagenesis of any of these residues compromised the integrity of the NrCAM-Npn2 interaction. Thus, the DDK network in NrCAM Ig1 and Npn2 Arg^31^ comprise a charged interface which, together with the NrCAM TARNER motif [10], promotes association of NrCAM and Npn2. Importantly, NrCAM stabilizes the Sema3F receptor complex by enhancing binding between Npn2 and PlexA3. Structural studies of a related receptor indicate that Sema3A induces receptor clustering and PlexA2 activation through Npn1 cross-bracing [16]. The relatively low affinity between Npn1 and PlexA2 (Kd 66 µM [16]) might increase in the presence of L1 or CHL1, as these L1CAMs mediate Sema3A-induced axon repulsion [6, 7].

We provide evidence here that Sema3F transduces intracellular signaling in apical dendritic spines through Tiam1-Rac1-PAK-LIMK-Cofilin and RhoA-ROCK-Myosin II, culminating in actin cytoskeletal remodeling and spine retraction. A postulated scheme for Sema3F signaling is depicted in Fig. 6. In accord with Sema3F-induced spine loss through Tiam1 and Rac1, spine retraction in response to ephrinA5/B2 and EphB receptors is also coupled to Tiam1 and Rac1 [57]. A role for RhoA-ROCK signaling in spine destabilization has been demonstrated in various vertebrate and invertebrate neurons (reviewed in [58]), as well as in traumatic brain injury [59]. An open question is how RhoA becomes activated upon Sema3F treatment (Fig. 6). One possibility is that inhibition of Rap1 by Sema3F [12] compromises activation of Rap-dependent RhoGAPs, such as ARAP3 or p190RhoGAP [60,61,58], favorable for RhoA activation. It should be noted that Rac1 and RhoA pathways have shown differential effects in promoting or inhibiting spine retraction [62-64,39]. These differences may be due to different ligands, sustained versus acute ligand stimulation [65], or developmental timing. Although our studies were conducted using cortical neuronal cultures, the demonstrated involvement of Rac1 and RhoA signaling pathways in spine pruning are consistent with *in vivo* mouse mutant phenotypes. It was recently reported that a photoactivable form of Rac1 targeted to the postsynaptic density of pyramidal cell synapses increased spine elimination in PFC circuits [66], and elicited spine shrinkage in primary motor cortex while causing erasure of synaptic memory traces [67]. Furthermore, constitutively active RhoA decreased spine density in cortical and hippocampal brain slices [68].

The multiplicity of substrates for downstream effectors suggests that additional intermediates not identified here may contribute to Sema3F signaling. For example, Tiam1-Rac1 could exert added effects by engaging the actin nucleation complex WAVE1-Arp2/3 to promote actin assembly [35]. As shown in hippocampal spines, PAKs can also activate MEK-ERK signaling, and directly phosphorylate MLC at Ser19 [69]. In addition to phosphorylation of MLC II, ROCKs indirectly regulate MLC by inactivating MLC phosphatase [70]. ROCK1/2 also phosphorylates LIMK, PTEN, and the microtubule effector CRMP2 [59]. A role for microtubules is suggested by a recent report that CRMP2 mediates Sema3F dendritic spine pruning in the dentate gyrus [71].

It is well-established in different cell contexts that Rac1 signaling through PAK-LIMK-Cofilin promotes the assembly of actin filaments [38, 72], whereas RhoA signaling though ROCK-Myosin II promotes actomyosin contraction [73]. Inhibition of Sema3F-induced spine retraction by Latrunculin A and Jasplakinolide support a role for both actin assembly and disassembly in spine elimination. An interesting interpretation is that Sema3F-induced RhoA signaling generates contractile force in spines that exerts tension on actin filaments assembled through Rac1 signaling. Actin filaments are known to transmit mechanical force generated by Myosin II, which can result in actin depolymerization [53,54,49]. For instance, in nerve growth cones, high levels of mechanical tension from actomyosin contraction enhance turnover of treadmilling F-actin, leading to a catastrophic loss of actin filaments [74].

Our studies may have broader impact for understanding how other repellent ligands (Sema5A, Ephrins) and cell adhesion molecules (SynCAM, ALCAM, Contactins, Cadherins) regulate dendritic spines and synapses critical for cortical circuitry. These molecules likely coordinate with other cellular responses to achieve synaptic elimination, including astrocyte-mediated phagocytosis through engulfment receptors (MerTK, MEGF10), microglia-mediated pruning (C1q, C3, CX3CR1) [75–77], and autophagy through Caspase 3 [77] and mTOR [78]. Moreover, our studies defining molecular signaling pathways for Sema3F-induced spine pruning may provide insight into mis-regulated mechanisms that lead to spine dysgenesis in diseases such as autism, schizophrenia, and bipolar disorder [3].

## Supporting information

Supplemental Figure 1 - ANOVA comparison

Supplemental Table 1 - Morphology Analysis

Supplemental Figure 2 - Tiam1 Co-IP repeats

## AVAILABILITY OF DATA

All data contained in this manuscript will be made available upon request.

## ETHICS APPROVAL

All animals used in this study were maintained in the University of North Carolina Animal Facility according to Institutional Animal Care and Use Committee policies in accordance with National Institutes of Health guidelines (UNC Protocol 18-073, approved February 28, 2018).

## CONSENT TO PARTICIPATE

Not applicable

## CODE AVAILABILITY

Not applicable.

## CONSENT FOR PUBLICATION

Not applicable.

## ACKNOWLEDGEMENTS

We acknowledge Dr. Chelsea Sullivan for analysis of Rho GTPases and Dr. Kelsey Murphy, Teva Smith, and Micah Council for assistance with experiments. We are grateful to Dr. Pablo Ariel, Director of the Microscopy Services Laboratory in the UNC Department of Pathology and Laboratory Medicine for expert advice and assistance with imaging analysis (supported by P30 CA016086 Cancer Center Core Grant to UNC Lineberger Comprehensive Cancer Center).

---

**Supplementary Table 1.** Multinomial regression analysis tested the effect of inhibitor treatment on spine morphology as described in the Results section, yielding p-values for significant differences in the log-odds of spines to be mushroom instead of thin or stubby in Fc-treated control neurons. At least 80 spines were scored for each condition.

**Supplementary Figure 1.** Scatter plots of spine density on individual neurons in the presence and absence of inhibitors in cultures treated with Fc (blue) or Sema3F-Fc (pink), used in 2-way ANOVA statistical analysis. Blue and pink lines connect the means of each group distribution.

**Supplementary Figure 2.** Replicate examples (A, B) of co-immunoprecipitation of Tiam1 and NrCAM from postnatal mouse synaptoneurosomes, shown by immunoprecipitation (IP) with Tiam1 antibodies or normal Ig (NIg), followed by immunoblotting (IB) for Tiam1 (near 250 kDa marker) and reprobing blots for NrCAM (130 kDa). Input lysates are shown on the right.

## References

1. Alvarez VA, Sabatini BL (2007) Anatomical and physiological plasticity of dendritic spines. Annu Rev Neurosci 30:79–97. doi:10.1146/annurev.neuro.30.051606.094222

2. Stein IS, Zito K (2019) Dendritic Spine Elimination: Molecular Mechanisms and Implications. Neuroscientist 25 (1):27–47. doi:10.1177/1073858418769644

3. Forrest MP, Parnell E, Penzes P (2018) Dendritic structural plasticity and neuropsychiatric disease. Nat Rev Neurosci 19 (4):215–234. doi:10.1038/nrn.2018.16

4. Duncan BW, Murphy KE, Maness P (2021) Molecular Mechanisms of L1 and NCAM Adhesion Molecules in Synaptic Pruning, Plasticity, and Stabilization. Frontiers in Cell and Developmental Biology. doi:https://doi.org/10.3389/fcell.2021.625340

5. Castellani V, Chedotal A, Schachner M, Faivre-Sarrailh C, Rougon G (2000) Analysis of the L1-deficient mouse phenotype reveals cross-talk between Sema3A and L1 signaling pathways in axonal guidance. Neuron 27 (2):237–249. doi:10.1016/s0896-6273(00)00033-7

6. Wright AG, Demyanenko GP, Powell A, Schachner M, Enriquez-Barreto L, Tran TS, Polleux F, Maness PF (2007) Close homolog of L1 and neuropilin 1 mediate guidance of thalamocortical axons at the ventral telencephalon. J Neurosci 27 (50):13667–13679

7. Bechara A, Nawabi H, Moret F, Yaron A, Weaver E, Bozon M, Abouzid K, Guan JL, Tessier-Lavigne M, Lemmon V, Castellani V (2008) FAK-MAPK-dependent adhesion disassembly downstream of L1 contributes to semaphorin3A-induced collapse. EMBO J 27 (11):1549–1562

8. Koropouli E, Kolodkin AL (2014) Semaphorins and the dynamic regulation of synapse assembly, refinement, and function. Curr Opin Neurobiol 27:1–7. doi:10.1016/j.conb.2014.02.005

9. Tran TS, Rubio ME, Clem RL, Johnson D, Case L, Tessier-Lavigne M, Huganir RL, Ginty DD, Kolodkin AL (2009) Secreted semaphorins control spine distribution and morphogenesis in the postnatal CNS. Nature 462 (7276):1065–1069. doi:10.1038/nature08628

10. Demyanenko GP, Mohan V, Zhang X, Brennaman LH, Dharbal KE, Tran TS, Manis PB, Maness PF (2014) Neural Cell Adhesion Molecule NrCAM Regulates Semaphorin 3F-Induced Dendritic Spine Remodeling. J Neurosci 34 (34):11274–11287. doi:10.1523/JNEUROSCI.1774-14.2014

11. Mohan V, Wade SD, Sullivan CS, Kasten MR, Sweetman C, Stewart R, Truong Y, Schachner M, Manis PB, Maness PF (2019) Close Homolog of L1 Regulates Dendritic Spine Density in the Mouse Cerebral Cortex Through Semaphorin 3B. J Neurosci 39 (32):6233–6250. doi:10.1523/JNEUROSCI.2984-18.2019

12. Mohan V, Sullivan CS, Guo J, Wade SD, Majumder S, Agarwal A, Anton ES, Temple BS, Maness PF (2019) Temporal Regulation of Dendritic Spines Through NrCAM-Semaphorin3F Receptor Signaling in Developing Cortical Pyramidal Neurons. Cereb Cortex 29 (3):963–977. doi:10.1093/cercor/bhy004

13. Gordon U, Polsky A, Schiller J (2006) Plasticity compartments in basal dendrites of neocortical pyramidal neurons. J Neurosci 26 (49):12717–12726. doi:10.1523/JNEUROSCI.3502-06.2006

14. Shigematsu N, Ueta Y, Mohamed AA, Hatada S, Fukuda T, Kubota Y, Kawaguchi Y (2016) Selective Thalamic Innervation of Rat Frontal Cortical Neurons. Cereb Cortex 26 (6):2689–2704. doi:10.1093/cercor/bhv124

15. Wang Q, Chiu SL, Koropouli E, Hong I, Mitchell S, Easwaran TP, Hamilton NR, Gustina AS, Zhu Q, Ginty DD, Huganir RL, Kolodkin AL (2017) Neuropilin-2/PlexinA3 Receptors Associate with GluA1 and Mediate Sema3F-Dependent Homeostatic Scaling in Cortical Neurons. Neuron 96 (5):1084–1098 e1087. doi:10.1016/j.neuron.2017.10.029

16. Janssen BJ, Malinauskas T, Weir GA, Cader MZ, Siebold C, Jones EY (2012) Neuropilins lock secreted semaphorins onto plexins in a ternary signaling complex. Nat Struct Mol Biol 19 (12):1293–1299. doi:nsmb.2416 [pii] 10.1038/nsmb.2416

17. Pascoe HG, Wang Y, Zhang X (2015) Structural mechanisms of plexin signaling. Prog Biophys Mol Biol 118 (3):161–168. doi:10.1016/j.pbiomolbio.2015.03.006

18. DePoy LM, Shapiro LP, Kietzman HW, Roman KM, Gourley SL (2019) beta1-Integrins in the Developing Orbitofrontal Cortex Are Necessary for Expectancy Updating in Mice. J Neurosci 39 (34):6644–6655. doi:10.1523/JNEUROSCI.3072-18.2019

19. Webb B, Sali A (2014) Comparative Protein Structure Modeling Using MODELLER. Curr Protoc Bioinformatics 47:5 6 1–32. doi:10.1002/0471250953.bi0506s47

20. Liu H, Focia PJ, He X (2011) Homophilic adhesion mechanism of neurofascin, a member of the L1 family of neural cell adhesion molecules. J Biol Chem 286 (1):797–805. doi:10.1074/jbc.M110.180281

21. Chen H, Chedotal A, He Z, Goodman CS, Tessier-Lavigne M (1997) Neuropilin-2, a novel member of the neuropilin family, is a high affinity receptor for the semaphorins Sema E and Sema IV but not Sema III [published erratum appears in Neuron 1997 Sep;19(3):559]. Neuron 19 (3):547–559

22. Comeau SR, Gatchell DW, Vajda S, Camacho CJ (2004) ClusPro: an automated docking and discrimination method for the prediction of protein complexes. Bioinformatics 20 (1):45–50

23. Kozakov D, Brenke R, Comeau SR, Vajda S (2006) PIPER: an FFT-based protein docking program with pairwise potentials. Proteins 65 (2):392–406. doi:10.1002/prot.21117

24. Duman JG, Tzeng CP, Tu YK, Munjal T, Schwechter B, Ho TS, Tolias KF (2013) The adhesion-GPCR BAI1 regulates synaptogenesis by controlling the recruitment of the Par3/Tiam1 polarity complex to synaptic sites. J Neurosci 33 (16):6964–6978. doi:10.1523/JNEUROSCI.3978-12.2013

25. Tolias KF, Bikoff JB, Kane CG, Tolias CS, Hu L, Greenberg ME (2007) The Rac1 guanine nucleotide exchange factor Tiam1 mediates EphB receptor-dependent dendritic spine development. Proc Natl Acad Sci U S A 104 (17):7265–7270

26. Hayashi-Takagi A, Araki Y, Nakamura M, Vollrath B, Duron SG, Yan Z, Kasai H, Huganir RL, Campbell DA, Sawa A (2014) PAKs inhibitors ameliorate schizophrenia-associated dendritic spine deterioration in vitro and in vivo during late adolescence. Proc Natl Acad Sci U S A 111 (17):6461–6466. doi:10.1073/pnas.1321109111

27. Garcia-Mata R, Wennerberg K, Arthur WT, Noren NK, Ellerbroek SM, Burridge K (2006) Analysis of activated GAPs and GEFs in cell lysates. Methods Enzymol 406:425–437. doi:10.1016/S0076-6879(06)06031-9

28. Villasana LE, Klann E, Tejada-Simon MV (2006) Rapid isolation of synaptoneurosomes and postsynaptic densities from adult mouse hippocampus. J Neurosci Methods 158 (1):30–36. doi:10.1016/j.jneumeth.2006.05.008

29. Mohan V, Gomez JR, Maness PF (2019) Expression and Function of Neuron-Glia-Related Cell Adhesion Molecule (NrCAM) in the Amygdalar Pathway. Front Cell Dev Biol 7:9. doi:10.3389/fcell.2019.00009

30. Peters A, Harriman KM (1990) Different kinds of axon terminals forming symmetric synapses with the cell bodies and initial axon segments of layer II/III pyramidal cells. I. Morphometric analysis. J Neurocytol 19 (2):154–174

31. Agresti A (2013) Categorical Data Analysis. third edn. Wiley, NY.,

32. Comeau SR, Gatchell DW, Vajda S, Camacho CJ (2004) ClusPro: a fully automated algorithm for protein-protein docking. Nucleic Acids Res 32 (Web Server issue):W96–99. doi:10.1093/nar/gkh354

33. Tran TS, Kolodkin AL, Bharadwaj R (2007) Semaphorin regulation of cellular morphology. Annu Rev Cell Dev Biol 23:263–292

34. Gao Y, Dickerson JB, Guo F, Zheng J, Zheng Y (2004) Rational design and characterization of a Rac GTPase-specific small molecule inhibitor. Proc Natl Acad Sci U S A 101 (20):7618–7623. doi:10.1073/pnas.0307512101

35. Boissier P, Huynh-Do U (2014) The guanine nucleotide exchange factor Tiam1: a Janus-faced molecule in cellular signaling. Cell Signal 26 (3):483–491. doi:10.1016/j.cellsig.2013.11.034

36. Dai J, Buhusi M, Demyanenko GP, Brennaman LH, Hruska M, Dalva MB, Maness PF (2013) Neuron glia-related cell adhesion molecule (NrCAM) promotes topographic retinocollicular mapping. PLoS One 8 (9):e73000. doi:10.1371/journal.pone.0073000

37. Bokoch GM (2003) Biology of the p21-activated kinases. Annu Rev Biochem 72:743–781

38. Civiero L, Greggio E (2018) PAKs in the brain: Function and dysfunction. Biochim Biophys Acta Mol Basis Dis 1864 (2):444–453. doi:10.1016/j.bbadis.2017.11.005

39. Hayashi ML, Choi SY, Rao BS, Jung HY, Lee HK, Zhang D, Chattarji S, Kirkwood A, Tonegawa S (2004) Altered cortical synaptic morphology and impaired memory consolidation in forebrain-specific dominant-negative PAK transgenic mice. Neuron 42 (5):773–787

40. Colgan LA, Yasuda R (2014) Plasticity of dendritic spines: subcompartmentalization of signaling. Annu Rev Physiol 76:365–385. doi:10.1146/annurev-physiol-021113-170400

41. Dolan BM, Duron SG, Campbell DA, Vollrath B, Shankaranarayana Rao BS, Ko HY, Lin GG, Govindarajan A, Choi SY, Tonegawa S (2013) Rescue of fragile X syndrome phenotypes in Fmr1 KO mice by the small-molecule PAK inhibitor FRAX486. Proc Natl Acad Sci U S A 110 (14):5671–5676. doi:10.1073/pnas.1219383110

42. Bernard O (2007) Lim kinases, regulators of actin dynamics. Int J Biochem Cell Biol 39 (6):1071–1076. doi:10.1016/j.biocel.2006.11.011

43. Cuberos H, Vallee B, Vourc’h P, Tastet J, Andres CR, Benedetti H (2015) Roles of LIM kinases in central nervous system function and dysfunction. FEBS Lett 589 (24 Pt B):3795–3806. doi:10.1016/j.febslet.2015.10.032

44. Mizuno K (2013) Signaling mechanisms and functional roles of cofilin phosphorylation and dephosphorylation. Cell Signal 25 (2):457–469. doi:10.1016/j.cellsig.2012.11.001

45. Yu Q, Gratzke C, Wang Y, Herlemann A, Sterr CM, Rutz B, Ciotkowska A, Wang X, Strittmatter F, Stief CG, Hennenberg M (2018) Inhibition of human prostate smooth muscle contraction by the LIM kinase inhibitors, SR7826 and LIMKi3. Br J Pharmacol 175 (11):2077–2096. doi:10.1111/bph.14201

46. Yin Y, Zheng K, Eid N, Howard S, Jeong JH, Yi F, Guo J, Park CM, Bibian M, Wu W, Hernandez P, Park H, Wu Y, Luo JL, LoGrasso PV, Feng Y (2015) Bis-aryl urea derivatives as potent and selective LIM kinase (Limk) inhibitors. J Med Chem 58 (4):1846–1861. doi:10.1021/jm501680m

47. Kanellos G, Frame MC (2016) Cellular functions of the ADF/cofilin family at a glance. J Cell Sci 129 (17):3211–3218. doi:10.1242/jcs.187849

48. Mueller BK, Mack H, Teusch N (2005) Rho kinase, a promising drug target for neurological disorders. Nat Rev Drug Discov 4 (5):387–398. doi:10.1038/nrd1719

49. Chugh P, Paluch EK (2018) The actin cortex at a glance. J Cell Sci 131 (14). doi:10.1242/jcs.186254

50. Uehata M, Ishizaki T, Satoh H, Ono T, Kawahara T, Morishita T, Tamakawa H, Yamagami K, Inui J, Maekawa M, Narumiya S (1997) Calcium sensitization of smooth muscle mediated by a Rho-associated protein kinase in hypertension [see comments]. Nature 389 (6654):990–994

51. Hodges JL, Newell-Litwa K, Asmussen H, Vicente-Manzanares M, Horwitz AR (2011) Myosin IIb activity and phosphorylation status determines dendritic spine and post-synaptic density morphology. PLoS One 6 (8):e24149. doi:10.1371/journal.pone.0024149

52. Ryu J, Liu L, Wong TP, Wu DC, Burette A, Weinberg R, Wang YT, Sheng M (2006) A critical role for myosin IIb in dendritic spine morphology and synaptic function. Neuron 49 (2):175–182. doi:10.1016/j.neuron.2005.12.017

53. Romet-Lemonne G, Jegou A (2013) Mechanotransduction down to individual actin filaments. Eur J Cell Biol 92 (10-11):333–338. doi:10.1016/j.ejcb.2013.10.011

54. Haviv L, Gillo D, Backouche F, Bernheim-Groswasser A (2008) A cytoskeletal demolition worker: myosin II acts as an actin depolymerization agent. J Mol Biol 375 (2):325–330. doi:10.1016/j.jmb.2007.09.066

55. Frost NA, Shroff H, Kong H, Betzig E, Blanpied TA (2010) Single-molecule discrimination of discrete perisynaptic and distributed sites of actin filament assembly within dendritic spines. Neuron 67 (1):86–99. doi:10.1016/j.neuron.2010.05.026

56. Gu J, Lee CW, Fan Y, Komlos D, Tang X, Sun C, Yu K, Hartzell HC, Chen G, Bamburg JR, Zheng JQ (2010) ADF/cofilin-mediated actin dynamics regulate AMPA receptor trafficking during synaptic plasticity. Nat Neurosci 13 (10):1208–1215. doi:10.1038/nn.2634

57. Um K, Niu S, Duman JG, Cheng JX, Tu YK, Schwechter B, Liu F, Hiles L, Narayanan AS, Ash RT, Mulherkar S, Alpadi K, Smirnakis SM, Tolias KF (2014) Dynamic control of excitatory synapse development by a Rac1 GEF/GAP regulatory complex. Dev Cell 29 (6):701–715. doi:10.1016/j.devcel.2014.05.011

58. Koleske AJ (2013) Molecular mechanisms of dendrite stability. Nat Rev Neurosci 14 (8):536–550. doi:nrn3486 [pii] 10.1038/nrn3486

59. Mulherkar S, Tolias KF (2020) RhoA-ROCK Signaling as a Therapeutic Target in Traumatic Brain Injury. Cells 9 (1). doi:10.3390/cells9010245

60. Jeon CY, Moon MY, Kim JH, Kim HJ, Kim JG, Li Y, Jin JK, Kim PH, Kim HC, Meier KE, Kim YS, Park JB (2012) Control of neurite outgrowth by RhoA inactivation. J Neurochem 120 (5):684–698. doi:10.1111/j.1471-4159.2011.07564.x

61. Artamonov MV, Jin L, Franke AS, Momotani K, Ho R, Dong XR, Majesky MW, Somlyo AV (2015) Signaling pathways that control rho kinase activity maintain the embryonic epicardial progenitor state. J Biol Chem 290 (16):10353–10367. doi:10.1074/jbc.M114.613190

62. Festa LK, Irollo E, Platt BJ, Tian Y, Floresco S, Meucci O (2020) CXCL12-induced rescue of cortical dendritic spines and cognitive flexibility. Elife 9. doi:10.7554/eLife.49717

63. Shapiro LP, Kietzman HW, Guo J, Rainnie DG, Gourley SL (2019) Rho-kinase inhibition has antidepressant-like efficacy and expedites dendritic spine pruning in adolescent mice. Neurobiol Dis 124:520–530. doi:10.1016/j.nbd.2018.12.015

64. Zhou L, Jones EV, Murai KK (2012) EphA signaling promotes actin-based dendritic spine remodeling through slingshot phosphatase. J Biol Chem 287 (12):9346–9359. doi:10.1074/jbc.M111.302802

65. Mao YT, Zhu JX, Hanamura K, Iurilli G, Datta SR, Dalva MB (2018) Filopodia Conduct Target Selection in Cortical Neurons Using Differences in Signal Kinetics of a Single Kinase. Neuron 98 (4):767–782 e768. doi:10.1016/j.neuron.2018.04.011

66. Moda-Sava RN, Murdock MH, Parekh PK, Fetcho RN, Huang BS, Huynh TN, Witztum J, Shaver DC, Rosenthal DL, Alway EJ, Lopez K, Meng Y, Nellissen L, Grosenick L, Milner TA, Deisseroth K, Bito H, Kasai H, Liston C (2019) Sustained rescue of prefrontal circuit dysfunction by antidepressant-induced spine formation. Science 364 (6436). doi:10.1126/science.aat8078

67. Hayashi-Takagi A, Yagishita S, Nakamura M, Shirai F, Wu YI, Loshbaugh AL, Kuhlman B, Hahn KM, Kasai H (2015) Labelling and optical erasure of synaptic memory traces in the motor cortex. Nature 525 (7569):333–338. doi:10.1038/nature15257

68. Tashiro A, Minden A, Yuste R (2000) Regulation of dendritic spine morphology by the rho family of small GTPases: antagonistic roles of Rac and Rho. Cereb Cortex 10 (10):927–938.

69. Zhang H, Webb DJ, Asmussen H, Niu S, Horwitz AF (2005) A GIT1/PIX/Rac/PAK signaling module regulates spine morphogenesis and synapse formation through MLC. J Neurosci 25 (13):3379–3388. doi:10.1523/JNEUROSCI.3553-04.2005

70. Kimura K, Ito M, Amano M, Chihara K, Fukata Y, Nakafuku M, Yamamori B, Feng J, Nakano T, Okawa K, Iwamatsu A, Kaibuchi K (1996) Regulation of myosin phosphatase by Rho and Rho-associated kinase (Rho-kinase). Science 273 (5272):245–248. doi:10.1126/science.273.5272.245

71. Ziak J, Weissova R, Jerabkova K, Janikova M, Maimon R, Petrasek T, Pukajova B, Kleisnerova M, Wang M, Brill MS, Kasparek P, Zhou X, Alvarez-Bolado G, Sedlacek R, Misgeld T, Stuchlik A, Perlson E, Balastik M (2020) CRMP2 mediates Sema3F-dependent axon pruning and dendritic spine remodeling. EMBO Rep 21 (3):e48512. doi:10.15252/embr.201948512

72. Spence EF, Soderling SH (2015) Actin Out: Regulation of the Synaptic Cytoskeleton. J Biol Chem 290 (48):28613–28622. doi:10.1074/jbc.R115.655118

73. Newell-Litwa KA, Horwitz R, Lamers ML (2015) Non-muscle myosin II in disease: mechanisms and therapeutic opportunities. Dis Model Mech 8 (12):1495–1515. doi:10.1242/dmm.022103

74. Zhang XF, Ajeti V, Tsai N, Fereydooni A, Burns W, Murrell M, De La Cruz EM, Forscher P (2019) Regulation of axon growth by myosin II-dependent mechanocatalysis of cofilin activity. J Cell Biol 218 (7):2329–2349. doi:10.1083/jcb.201810054

75. Weinhard L, di Bartolomei G, Bolasco G, Machado P, Schieber NL, Neniskyte U, Exiga M, Vadisiute A, Raggioli A, Schertel A, Schwab Y, Gross CT (2018) Microglia remodel synapses by presynaptic trogocytosis and spine head filopodia induction. Nat Commun 9 (1):1228. doi:10.1038/s41467-018-03566-5

76. Wilton DK, Dissing-Olesen L, Stevens B (2019) Neuron-Glia Signaling in Synapse Elimination. Annu Rev Neurosci 42:107–127. doi:10.1146/annurev-neuro-070918-050306

77. Erturk A, Wang Y, Sheng M (2014) Local pruning of dendrites and spines by caspase-3-dependent and proteasome-limited mechanisms. J Neurosci 34 (5):1672–1688. doi:10.1523/JNEUROSCI.3121-13.2014

78. Lieberman OJ, McGuirt AF, Tang G, Sulzer D (2019) Roles for neuronal and glial autophagy in synaptic pruning during development. Neurobiol Dis 122:49–63. doi:10.1016/j.nbd.2018.04.017

