## Supplementary figures and images for "Semaphorin3F Drives Dendritic Spine Pruning through Rho-GTPase Signaling"

### Supplemental Figure 1 - ANOVA comparison

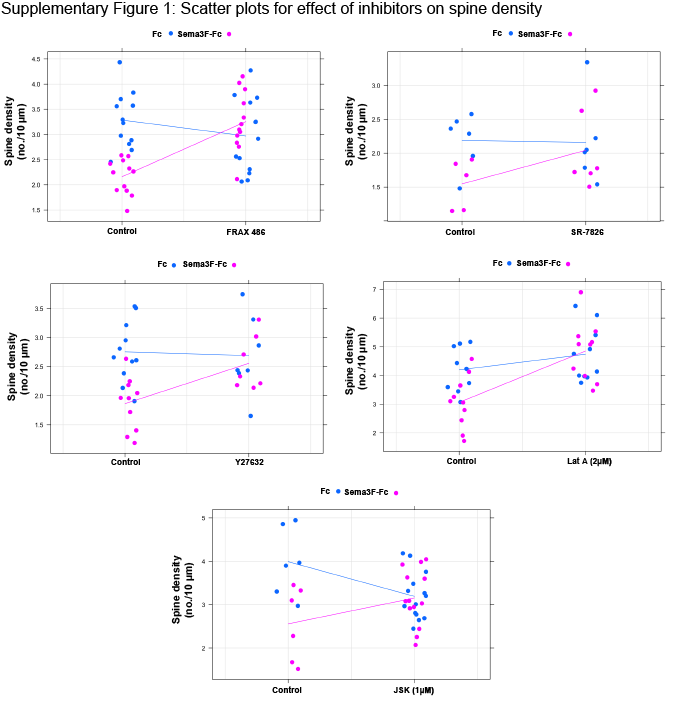

### Supplemental Figure 2 - Tiam1 Co-IP repeats

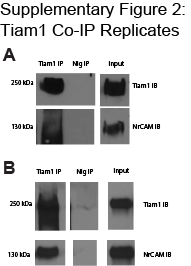

### Supplemental Table 1 - Morphology Analysis

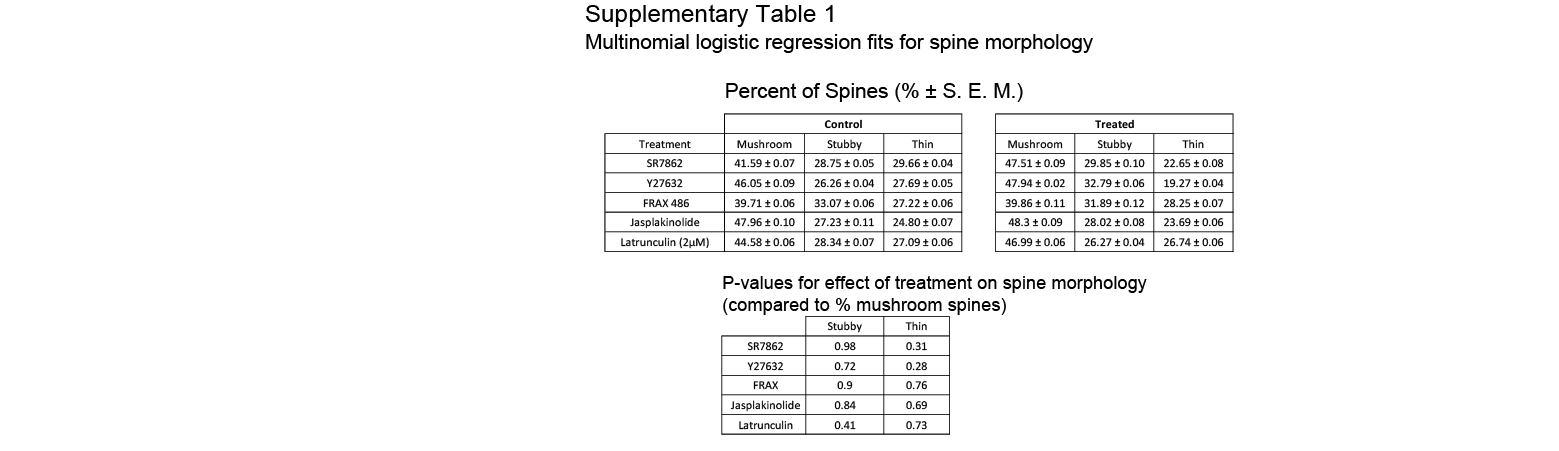
